# Individual differences drive social hierarchies in male mouse societies

**DOI:** 10.1101/2025.08.26.672308

**Authors:** Jonathan R. Reinwald, Sarah Ghanayem, David Wolf, Max F. Scheller, Julia Lebedeva, Philipp Lebhardt, Oliver Gölz, Corentin Nelias, Wolfgang Kelsch

**Affiliations:** Dept. of Psychiatry and Psychotherapy, University Medical Center Mainz, Johannes-Gutenberg University, Untere Zahlbacher Strasse 8, 55131 Mainz, Germany; Dept. of Psychiatry and Psychotherapy, Central Institute of Mental Health, Medical Faculty Mannheim, Heidelberg University, Square J5, 68159 Mannheim, Germany; Dept. of Neuroimaging, Central Institute of Mental Health, Medical Faculty Mannheim, Heidelberg University, Square J5, 68159 Mannheim, Germany; Dept. of Clinical Health Technologies, Institute for Manufacturing Engineering and Automation, Fraunhofer Society, Theodor-Kutzer-Ufer 1-3, 68167 Mannheim, Germany

**Keywords:** social hierarchy, social status, naturalistic housing, chasing, dominance, subordination, non-shared environment, behavioral traits, trait stability, reinforcement learning, impulsivity, individual differences, individuality, mouse societies, mouse personalities, resilience

## Abstract

Social hierarchies structure groups and confer advantages on high-ranking individuals. In mice, individual position in hierarchies may emerge situationally from current group compositions, or, alternatively, may remain largely stable across groups as an internalized feature. Dominance and subordination are expressed in behaviors like tube competitions or agonistic chasing. The interaction of these behaviors in the shaping of social position in larger male mouse groups remains largely unknown. To address these questions, we developed the NoSeMaze, a semi-naturalistic, open-source, modular platform that enables automated long-term tracking of unperturbed groups. Across more than 4,000 mouse-days, hierarchies derived from incidental competitions in the integrated tube tests were non-despotic, transitive, and stable even when group compositions changed. This stability supports an internalized component of competition-based social rank. Chasing was also stable across contexts. Notably, chasing was concentrated among high-ranking individuals, consistent with ongoing negotiation of social rank among individuals at the upper end of the hierarchy. The link between chasing and social rank strengthened in groups with less well-defined rank structure, where mice rely more on aggressive signaling to assert their position. Chasing and social rank were associated with certain dimensions of simultaneously measured physical and cognitive features. In summary, high-dimensional tracking with the NoSeMaze reveals that social position in mice is multifaceted and shaped by stable dimensions of individual behavior that persist across changing social contexts. The approach thus enables longitudinal modeling of individuality and social position as key resilience factors.

## INTRODUCTION

In many social species, individuals organize into hierarchical structures characterized by dominance and subordination relative to others. Social rank shapes access to resources and reduces conflict ^1–3^, and is the most prominent yet identified resilience factor ^4–6^. In mice, social position is expressed across different behavioral domains, including competitive encounters and agonistic behaviors such as chasing ^7–9^. However, the precise nature of the relationship between these behavioral domains remains unclear, particularly in relatively stable social groups and under semi-naturalistic longitudinal conditions ^9–13^. Moreover, prior studies have yielded inconsistent findings on whether specific facets of social position are associated with individual differences in reward-related behaviors, such as impulsivity, cognitive flexibility, and inhibitory control ^14–16^.

Social structures in animals are often assumed to emerge from group dynamics that unfold over time ^2,3^. These processes are difficult to capture with reductionist paradigms ^17–20^ that are not designed for tracking moment-to-moment interactions over long periods and under ecologically valid conditions. Classic assays typically assess social and cognitive behaviors separately, often in different contexts or at different time points, limiting our ability to test how these domains interact and how stable they are within individuals ^17^. Moreover, assays such as the tube test ^21^ rely on forced encounters and manual handling, which may introduce confounds ^22,23^ and thereby distort natural behavior.

In mice, small groups tend to form highly despotic hierarchies ^24–26^, whereas larger groups exhibit more complex structures ^27^. These patterns suggest a strong influence of emergent group-level dynamics on social structure ^12,27^. Yet, animals do not enter social groups as blank slates ^28–30^; stable latent factors in the individual may also contribute to hierarchy formation ^13^. Thus, it remains unclear to what extent an individual’s social position is internalized and persists across different social contexts ^31^, or instead is primarily an emergent property of group-level dynamics. Disentangling these possibilities requires experimental conditions that allow unperturbed, continuous tracking of all individuals in sufficiently large groups, together with systematic changes of group composition to modulate social context.

While stable hierarchies and behavioral identity domains have been described previously in semi-naturalistic settings ^9–13^, many studies quantify these features either within fixed group compositions or in separate assays. Here, we therefore use systematic group reshuffling in 10-member societies to directly test whether individual differences in social rank and chasing persist across distinct social groups. In this study, we use social hierarchy to denote the group-level structure inferred from incidental competitions in the integrated tube tests, and social rank for an individual’s level within that hierarchy. Social position serves as an umbrella term for social rank and chasing behaviors. By continuously measuring social rank, chasing, and reinforcement-learning behavior in the same individuals, we further test how chasing and social rank are related to each other and how they relate to individual styles in non-social reinforcement learning.

To enable these tests, we developed the Non-invasive Sensor-rich Maze (NoSeMaze). The NoSeMaze provides a semi-naturalistic environment for fully automated, continuous, long-term behavioral assessment of individual mice living in groups of ten. Importantly, the system continuously monitors social and cognitive domains in parallel, without human intervention. Specifically, the system tracks social rank as real-time outcomes of incidental competitions in integrated tube tests, chasing, and self-initiated reinforcement learning via a go/no-go olfactory task. By combining these measurements across multiple group compositions, the NoSeMaze enables testing of the relative contributions of stable individual characteristics to social rank and chasing, and of how social and cognitive domains interact in shaping an individual’s social position.

The resulting longitudinal dataset, encompassing more than 4,000 mouse-days, was used to address three specific goals: (1) establish a high-resolution, longitudinal framework that captures individualized profiles of competition-based social rank, chasing, and reinforcement learning in group-living mice; (2) disentangle how chasing relates to hierarchy structure derived from incidental competitions in the integrated tube tests, and whether either behavior associates with individual differences in non-social reward-seeking; and (3) determine whether social rank, chasing, and non-social reward-seeking behaviors represent stable individual characteristics or dynamic features across time and changing group composition. As a secondary analysis, motivated by oxytocin’s established role in social recognition memory ^32,33^, we also tested whether OXTR deletion in the anterior olfactory nucleus produces detectable shifts in rank dynamics.

## Results

### Ecological longitudinal assessment in the NoSeMaze

The NoSeMaze provides a complex, semi-naturalistic environment designed for long-term automated assessment of reinforcement learning and social-position-related behaviors in mouse societies. It contains an open-field arena and a housing arena with nesting material and free access to food (**Fig. 1A, Supplementary Fig. S1**). The arenas are connected by two tubes equipped with RFID readers, which enable automated tracking of social interactions by detecting the identity and movement of individual RFID-tagged mice. Access to water is provided via the olfactory stimulus-outcome learning module, in which mice perform trials ad libitum to meet their water needs. The module is connected to the open-field arena, requiring mice to cross the tubes regularly to access food and water.

**Fig. 1:**
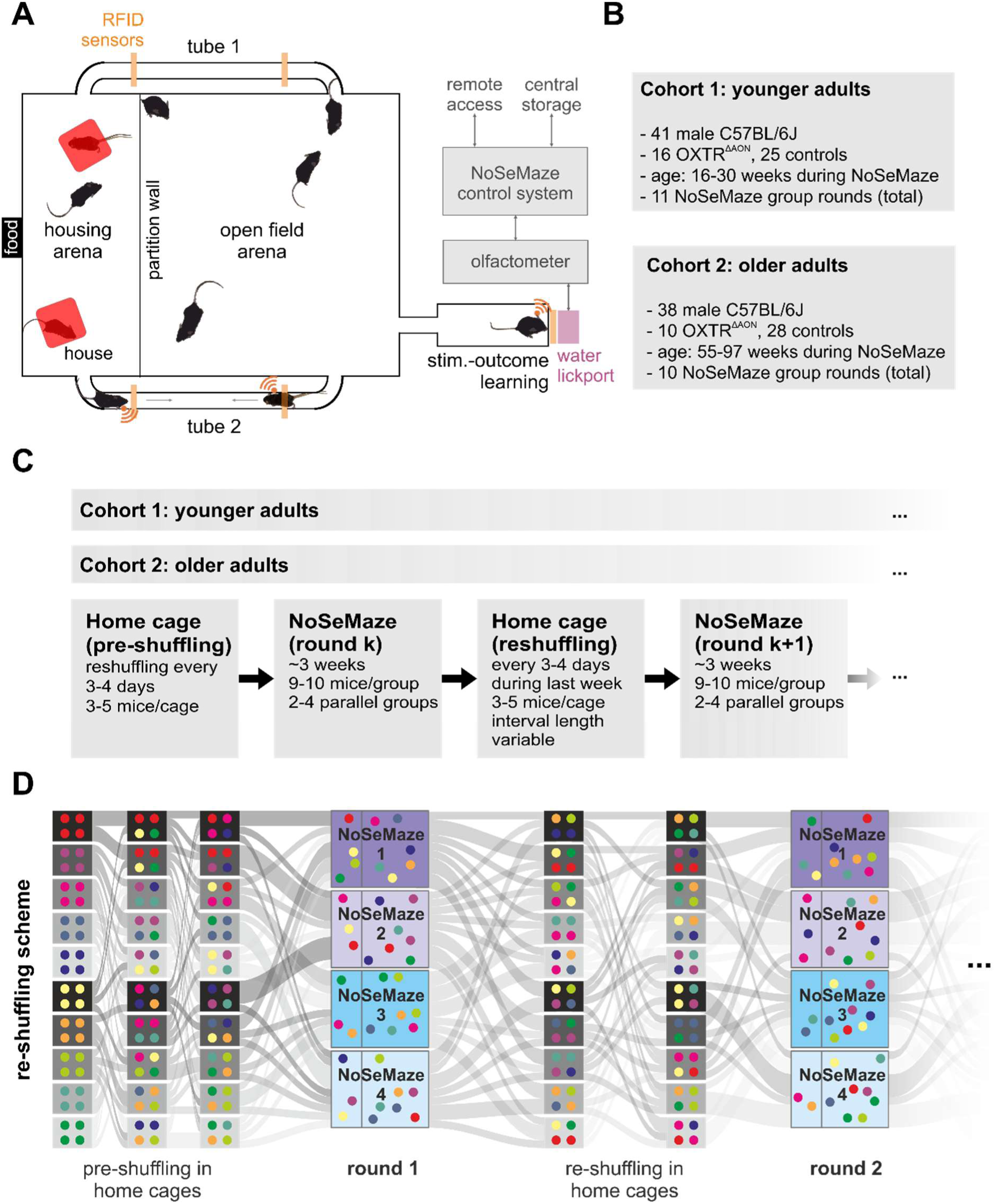
NoSeMaze setup, cohort structure, and accelerated longitudinal reshuffling design. **A**, The NoSeMaze (see **Supplementary Fig. S1**) tracks social rank, chasing, and reinforcement learning in mouse societies (n = 9-10 per group) over multiple weeks without human intervention. RFID-tagged animals are automatically identified at the tube entrances and during olfactory stimulus-outcome learning tasks at the water lickport. **B**, Overview of the two adult cohorts studied in separate experimental series. Cohort 1 (“younger adults”) comprised 41 male C57BL/6J-background mice (16 OXTR^ΔAON^, 25 controls), aged 16–30 weeks during NoSeMaze testing, and contributed 11 NoSeMaze group rounds in total. Cohort 2 (“older adults”) comprised 38 male mice (10 OXTR^ΔAON^, 28 controls), aged 55–97 weeks during NoSeMaze testing, and contributed 10 NoSeMaze group rounds in total. **C**, Experimental timeline. Before entry into the NoSeMaze, home-cage compositions were repeatedly reshuffled every 3–4 days (3–5 mice per cage) to promote familiarity among cohort members. Mice were then housed in the NoSeMaze for approximately 3 weeks per round in groups of 9–10, with 2–4 groups run in parallel. Between consecutive NoSeMaze rounds, mice returned to reshuffled home cages, where they were again reshuffled in the last week before the next NoSeMaze round. Reshuffling occurred within cohorts only, not across cohorts. **D**, Example of the reshuffling design across two consecutive NoSeMaze rounds. Colored dots represent individual mice and gray ribbons indicate reassignment from pre-shuffling home cages to NoSeMaze round 1, then to reshuffled home cages, and finally to NoSeMaze round 2. This accelerated longitudinal design systematically altered social group composition across rounds while maintaining animals within their age-defined cohort. *NoSeMaze, Non-invasive Sensor-rich Maze; RFID, radio frequency identification*

We investigated a large population of 79 male mice on the C57BL/6J background that were homozygous for floxed oxytocin receptor alleles (OXTRfl/fl), enabling conditional deletion of the receptor. In 26 mice, OXTR was selectively deleted in the AON pars centralis (OXTR^ΔAON^) via viral recombination, induced only after mice had reached full adulthood ^33^. OXTR^ΔAON^ mice show selective impairments in de-novo social olfactory recognition learning ^32,33^. The remaining 53 mice had normal OXTR expression. This manipulation was included as a secondary biological perturbation. The primary analyses and conclusions focus on the platform and cross-context stability, and genotype is treated as a covariate unless stated otherwise.

The NoSeMaze was originally optimized for male mice in which social hierarchies had been mostly characterized ^9^. Each mouse carried a subcutaneous RFID chip for identification. The study population comprised two age cohorts: younger (16-30 weeks) and older adults (55-97 weeks) (**Fig. 1B**). The two age cohorts were run as separate experimental series, and group reshuffling was performed within each cohort (**Fig. 1C**, see **Supplementary Table S2**). Mice lived in groups of 9-10 for multiple rounds in the NoSeMaze, with different group members in each round (**Fig. 1D**, see **Supplementary Table S1**). This allowed us to test which individual behaviors were stable across different group compositions. Before the start of the experiment, mice had been housed in groups of three to five in their home cages, with regular reshuffling to ensure familiarity among potential group members (**Fig. 1C-D**). To prevent bias in group assignment and behavioral interpretation, both the composition of experimental NoSeMaze groups and the home-cage groupings between rounds were assigned randomly and blinded to prior behavioral outcomes, individual performance, or genotype. Two to four NoSeMazes operated in parallel, with each round spanning three weeks. Between rounds, mice were reassigned to different home cage groups over several weeks (mean between round time: 3.8 weeks; SD: 2.8 weeks, see also **Supplementary Table S1**). In total, more than 4,000 mouse-days from 21 NoSeMaze rounds were analyzed (see **Supplementary Table S2**).

### Individual styles of reinforcement learning in the NoSeMaze

Before the first round in the NoSeMaze, mice were acclimatized to the reward port (see Methods). To assess individual differences in reinforcement learning, they then performed a go/no-go task with two olfactory conditioned stimuli (CS) in the stimulus-outcome learning module. Mice initiated trials by inserting their head into the odor port, where they were identified via RFID. In go trials, licking twice during the CS predicting reward (CS+) led to water delivery, while licking twice during the CS predicting no reward (CS−) in no-go trials triggered a timeout of six seconds (**Fig. 2A**). Reward contingencies alternated every three days between the two odors (i.e., ‘reversal’; **Fig. 2B**). As expected, activity levels peaked during the dark cycle (**Fig. 2C**). On average, mice performed 463 trials per day (**Supplementary Fig. S2A**). The number of trials required to reach 80% correct hits and correct rejections increased after the first reversal, suggesting that the initial reversal was challenging. With subsequent reversals, performance gradually improved, indicating that mice generally learned the task structure (**Fig. 2D**).

**Fig. 2:**
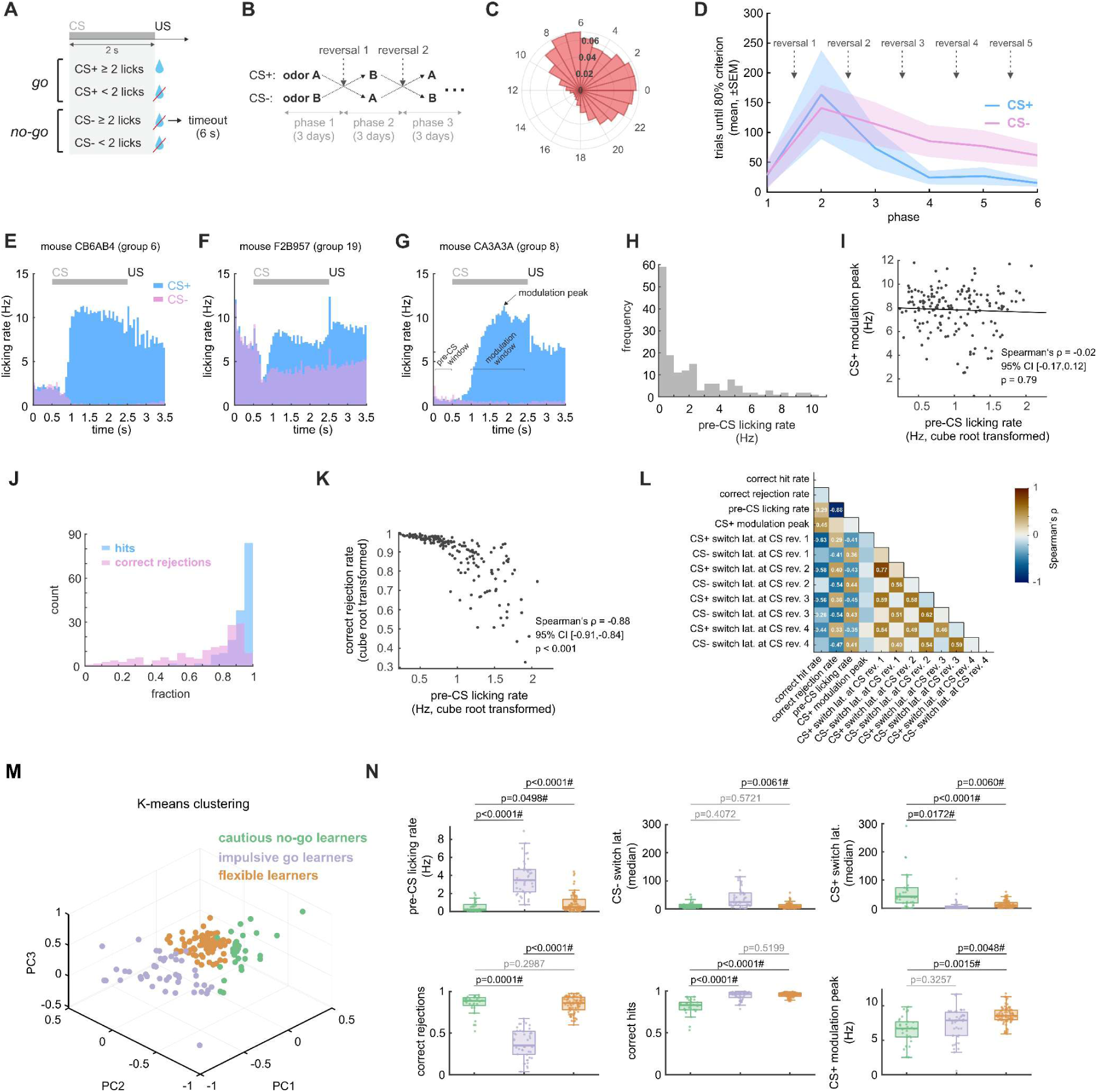
Inter-individual differences in reinforcement learning in the NoSeMaze. **A**, Mice performed stimulus-outcome learning trials for daily water intake, initiated by head insertion into the odor port. In CS+ trials, licking twice during odor presentation released water, while in CS− trials, it triggered a 6-second timeout. **B**, Reward contingencies switched every three days. **C**, Trial activity peaked during the dark phase. **D**, Number of trials required to reach 80% performance (hits or correct rejections, mean ± SEM) in each phase (cf. Fig. 2B). Performance decreased after the first reversal and improved in later phases, consistent with learning the task structure. Only data from the first round in the NoSeMaze was considered. **E-G**, Example lick patterns from three mice show variability in pre-CS licking rates and modulations during CS presentation in the stable phases of the last 150 trials before reversals. **H**, Pre-CS licking rates varied widely across animals. **I**, No correlation was observed between pre-CS licking rates and CS+ modulation peaks (see **Supplementary Fig. S2B**). **J**, The fraction of correct rejections was more broadly distributed across mice than hits. **K**, Pre-CS licking rates negatively correlated with correct rejection rates (see **Supplementary Fig. S2C**). **L**, Spearman’s correlation matrix (data from 17 groups) revealed significant relationships between reinforcement-learning metrics, including pre-CS licking rate, correct hit and rejection rates, CS+ modulation peak, and latency to switch after contingency reversals. Only statistically significant correlation coefficients are shown (Bonferroni-corrected for multiple comparisons). **M**, K-means clustering based on reward-seeking features identified three distinct learning strategies: impulsive go learners (purple), cautious no-go learners (green), and flexible learners (orange). **N**, The three learning strategies (color labeling as in **M**) differed across reward-seeking features including hit and rejection rates, pre-CS licking rates, CS switch latencies, and CS+ modulation peak. Horizontal bars denote statistical comparisons between clusters (p-values from permutation tests with n = 100,000 permutations using the median). P-values were corrected for multiple comparisons using the Benjamini-Hochberg false discovery rate (FDR) procedure (α = 0.05). Significant comparisons after FDR correction are marked with #. CI, confidence interval; CS+, rewarded conditioned stimulus; CS−, unrewarded conditioned stimulus; SEM, standard error of the mean Note: The regression line is illustrative only; statistical inference is based on Spearman’s correlation, which does not assume linearity.

Licking behavior varied widely across individuals, as shown for stable phases of the last 150 trials before reversals (**Fig. 2E-G**). Inter-individual differences were already seen in the pre-CS licking rates (**Fig. 2H**) that may serve as a proxy of impulsivity ^34^. Mice also differed in their licking patterns during reward anticipation. While all mice exhibited increased anticipatory licking during CS+ trials (CS+ modulation peak, cf. **Fig. 2E-G**), this measure of reward sensitivity was independent of pre-CS licking rates, suggesting distinct underlying mechanisms (**Fig. 2I**, *ρ = -*0.02, p = 0.79, **Supplementary Fig. S2B**). Correct hit rates were relatively high across animals (**Fig. 2J**), indicating effective CS+ recognition. The high variability in pre-CS licking and correct rejection rates (**Fig. 2J**) among individuals may again reflect individual differences in impulsivity and inhibitory control ^34^. Pre-CS licking was strongly negatively correlated with correct rejection rates (**Fig. 2K**, *ρ = -*0.88, p < 0.001, **Supplementary Fig. S2C**), suggesting impaired response suppression in these mice, leading to more timeouts in CS− trials (cf. **Fig. 2B**).

To quantify behavioral flexibility among mice, we measured CS+ and CS− switch latencies, defined as the trials required to restore (70% correct hits for CS+) or suppress (50% correct rejections for CS−) licking relative to each animal’s pre-reversal CS+ rate. As individualized measures, these latencies capture flexible adaptation rather than absolute performance. Higher pre-CS licking was associated with shorter CS+ switch latencies but slower adaptation to CS− reversals (**Fig. 2L**, see **Supplementary Material – Systematic Statistical Reporting** and **Supplementary Fig. S3**), suggesting that impulsivity facilitates detection of the rewarded cue but hinders economic response suppression.

We therefore examined whether multiple reward-seeking features cluster individuals into different learning strategies. K-means clustering based on CS+ and CS− median switch latencies, correct hit and rejection rates, CS+ modulation peak, and pre-CS licking identified three distinct learning strategies (**Fig. 2M**). *Cautious no-go learners* (**Fig. 2N**, green) exhibited the lowest impulsivity and adapted quickly to CS− reversals but updated CS+ associations more slowly, resulting in the lowest correct hit rates. In contrast, *impulsive go learners* (**Fig. 2N**, purple), characterized by the highest pre-CS licking, quickly adjusted to CS+ reversals but exhibited poor inhibitory control during CS− reversals, leading to frequent timeouts and a low correct rejection rate. Lastly, *flexible learners* (**Fig. 2N**, orange), who showed relatively low impulsivity, efficiently adapted to both CS+ and CS− reversals, achieving high performance in correct hits and rejections. This last cluster thus displayed an optimal balance between retrieving rewards with minimal timeouts while adapting effectively when cognitive flexibility was required. Flexible learners formed the largest group and also exhibited the strongest modulation of anticipatory CS+ licking.

Detailed statistical reporting for all analyses, including *n*, degrees of freedom, and 95% confidence intervals, is provided in the **Supplementary Material - Systematic Statistical Reporting**. **Extended Data** includes additional linear mixed-effects models controlling for potential confounding variables.

In summary, the NoSeMaze provides quantifiable measures of reinforcement learning, including flexibility, response adaptation, and impulsivity. The three behavioral clusters demonstrate how the NoSeMaze captures individual differences in learning flexibility and inhibitory control.

### Social interactions in the tubes

The NoSeMaze enables automated, real-time tracking of two distinct social interactions in the tubes: incidental dyadic competitions and voluntary tube chasing. Here, social hierarchy refers to the group-level structure reconstructed from incidental competitions in the integrated tube tests, and social rank to the individual’s level herein. Social position is used as an umbrella term for both social rank and chasing.

Tube competitions occur incidentally when two mice enter the tube from opposite ends at the same time, revealing dominance as the other retreats (**Fig. 3A**; see **Supplementary Videos S1–3** for insightful examples). The dominant mouse is detected sequentially at both ends, whereas the subordinate is detected twice at its entry site – once upon entry and again when retreating. Each dyadic tube competition results in symmetrical win/loss outcomes, enabling the construction of interaction networks and social ranks using David’s scores (**Fig. 3B**), which also consider the opponents’ social ranks. Competitions occurred primarily during the dark cycle (**Supplementary Fig. S4A-B**).

**Fig. 3:**
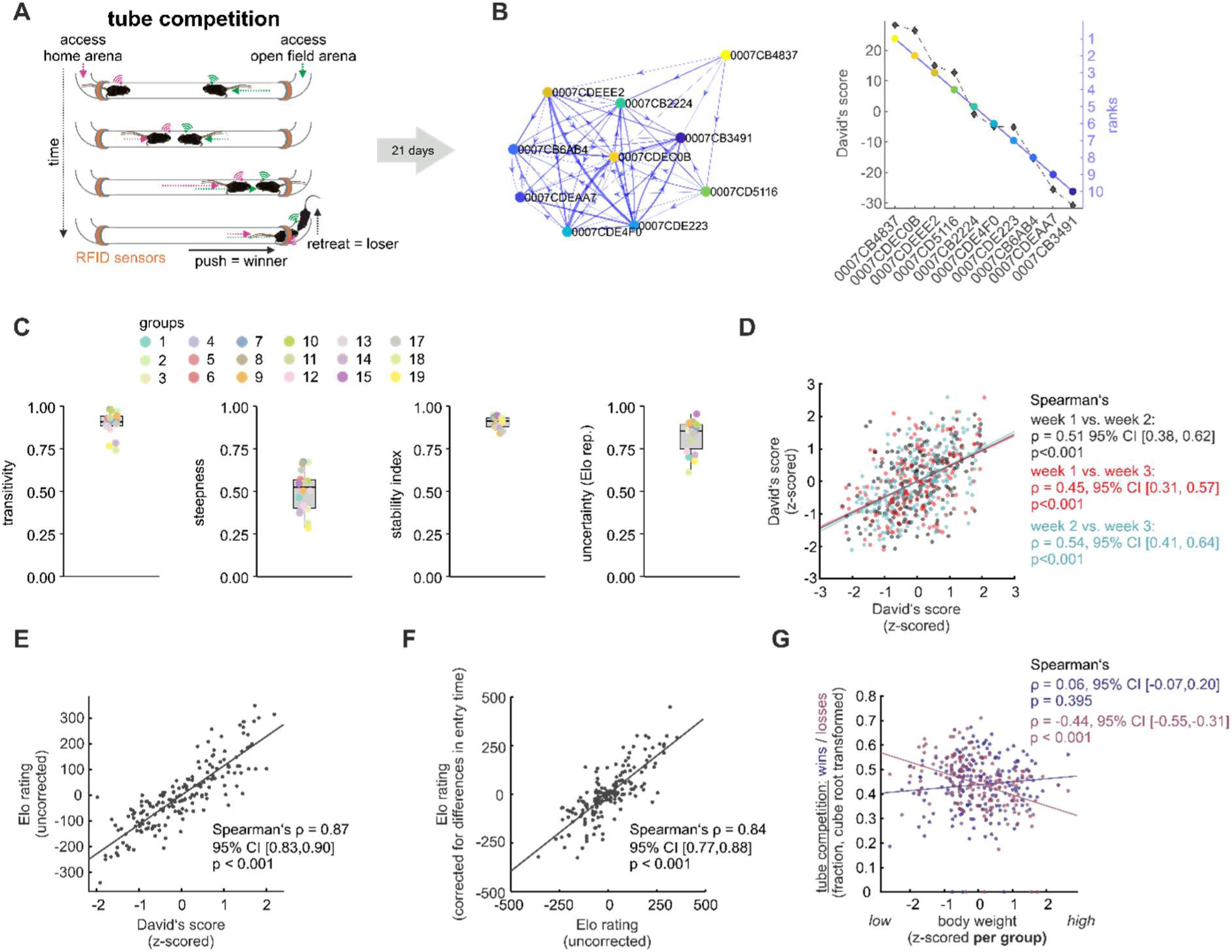
Social rank derived from incidental competitions in the integrated tube tests of the NoSeMaze. **A**, Incidental encounters in the tubes triggered dyadic tube competitions. **B**, Network graph derived from the cumulative tube competitions in an example group over three weeks. Outgoing arrows indicate wins, incoming arrows indicate losses. Individual hierarchical positions were calculated using the David’s score and can be converted into linear social ranks from 1 (highest) to 10 (lowest). **C**, Box plots of metrics characterizing social hierarchy for 18 groups, including transitivity, steepness, stability, and uncertainty-by-repeatability (for details, see ‘Source Data’). Group labels correspond to original experimental group IDs. Only groups with available tube-competition data are shown. Therefore, numbering is non-consecutive (e.g., groups 16, 20, and 21 are absent; see **Supplementary Table S2**). **D**, Scatterplots show that David’s scores (z-scored) were significantly correlated across the three different weeks, indicating temporal stability of social rank within one NoSeMaze round (data from 18 groups). **E**, David’s scores were strongly correlated with Elo ratings, demonstrating convergent validity of social rank measures. **F**, Elo ratings corrected for differences in entry time highly correlated to uncorrected Elo ratings. **G**, Relative body weight (z-scored per group) was significantly negatively correlated to the fraction of losses in tube competitions (cube root transformed, red), with heavier animals being less likely to lose. No association was found between body weight and the fraction of wins in tube competitions (cube root transformed, dark blue). Body weight was measured prior to the animals’ introduction to the NoSeMaze, respectively. CI, confidence interval; SD, standard deviation; Note: Regression lines are illustrative only; statistical inference is based on Spearman’s correlation, which does not assume linearity.

Social hierarchies derived from tube competitions were well-structured and stable, showing transitivity values exceeding 0.7 and moderate steepness across groups (**Fig. 3C**, see ‘Source Data’). Stability scores above 0.8 and high uncertainty-by-repeatability values ^35^ confirmed robust social hierarchy sampling over the three-week period (**Fig. 3C**). David’s scores also showed stable correlations across weeks (**Fig. 3D**, *ρ =* 0.45 – 0.54, all p < 0.001), indicating consistent social ranks over time. Moreover, David’s scores correlated strongly with dynamic social hierarchy measures such as the Elo rating (**Fig. 3E**, *ρ =* 0.87, p < 0.001), demonstrating their convergent validity in capturing social rank. Notably, social ranks with and without adjustment for entry time differences were highly correlated (**Fig. 3F**, *ρ =* 0.84, p < 0.001), supporting that social rank assessments were largely independent of entry time. Finally, David’s scores from the NoSeMaze significantly correlated with those from a manual tube test conducted after the experiment, when animals were again individually housed (**Supplementary Fig. S4C**).

Body weight showed a moderate positive association with social rank (**Supplementary Fig. S5A**). Interestingly, this association was driven by a significantly lower fraction of losses in heavier individuals (**Fig. 3G**, *ρ = -*0.44, p < 0.001, **Supplementary Fig. S5B**), while body weight did not correlate to the fraction of wins (**Fig. 3G**, *ρ =* 0.06, p = 0.395, **Supplementary Fig. S5C**). Thus, larger mice were less likely to be defeated in tube competitions, but did not win more frequently.

Unlike incidental tube competitions, tube chasing is a voluntarily initiated interaction in which one mouse actively chases another through the tube (**Fig. 4A**). To quantify these behaviors, we calculated the fraction of actively initiated chases and the fraction of times being chased for each mouse, relative to the total number of chasing events in its group (**Fig. 4A**, **Supplementary Fig. S6A**, left panel). Chasing occurred predominantly at night (**Fig. 4B**). The expression of chasing varied widely across individuals (cf. **Fig. 4A**, **Supplementary Fig. S6A**, left panel) and was notably skewed.

**Fig. 4:**
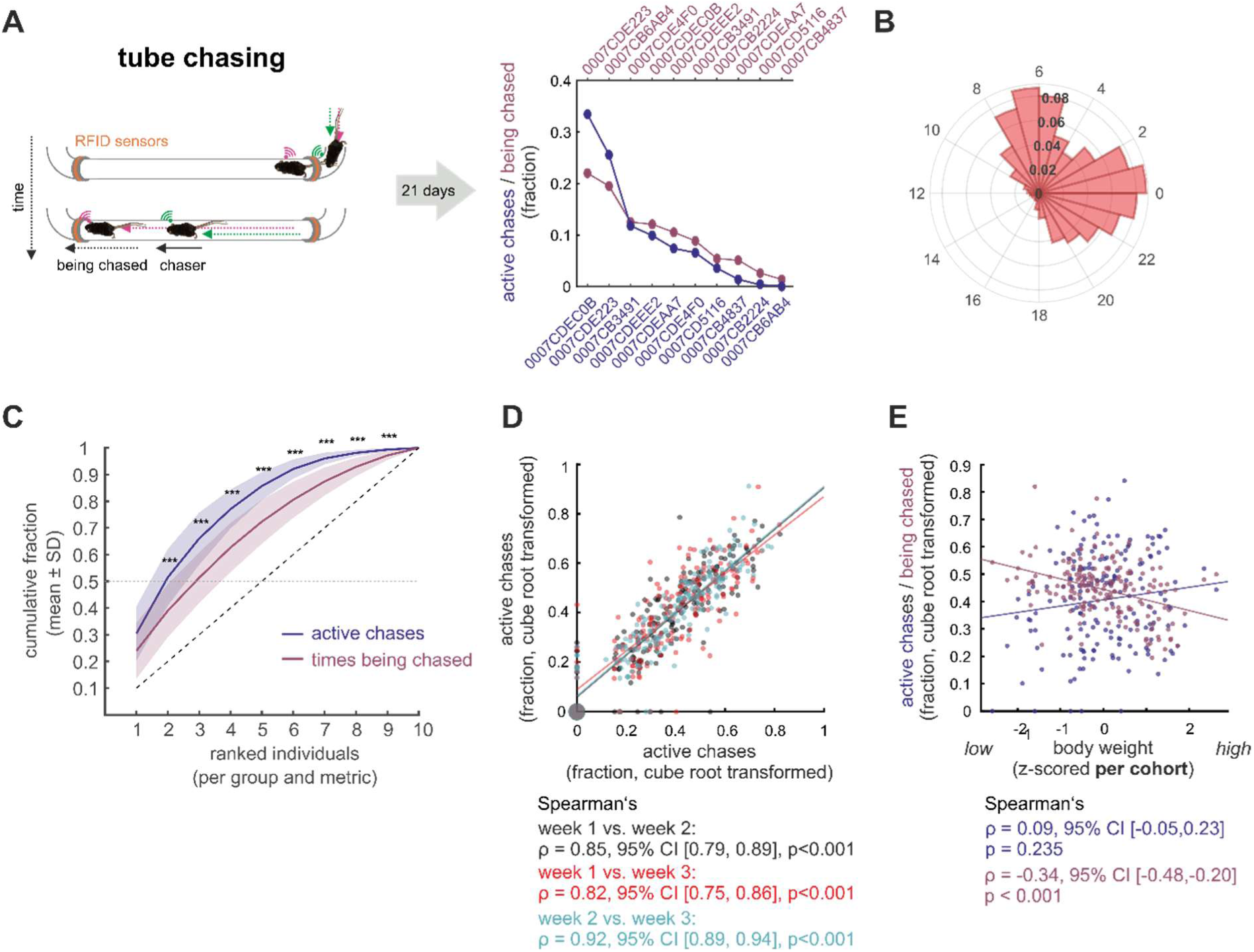
Chasing in the NoSeMaze. **A**, Schematic of the NoSeMaze setup for automated tracking of tube chasing. Chasing was quantified as the fraction of chases initiated by each individual relative to the total number of chases in their group (‘active chases’), as well as by the fraction of times being chased (see **Supplementary Fig. S6A**). **B**, Chasing occurred predominantly during the dark phase. **C**, Cumulative fraction of chases initiated (dark blue) and received (dark red) by individuals within each group, ordered by the fraction of chases they initiated. Chasing was concentrated in a small subset of individuals, with the two most active chasers accounting for more than 50% of all chases. Significant stars mark differences between the two curves (unpaired permutation test, n=10.000 permutations, ***, p < 0.001). **D**, Active chases were highly consistent across weeks. **E**, Active chases were not significantly correlated to relative weight before entering the NoSeMaze. However, the fraction of times being chased was negatively correlated with weight, indicating that heavier mice were less likely to be chased. CI, confidence interval; SD, standard deviation; Note: Regression lines are illustrative only; statistical inference is based on Spearman’s correlation, which does not assume linearity.

More specifically, a small subset of mice initiated most chases (**Fig. 4C**), with the two most active chasers accounting for more than 50% of all chases in a group. Across weeks, individual chasing behavior remained stable within the same NoSeMaze round (**Fig. 4D**, *ρ =* 0.82 to 0.92, all p < 0.001, **Supplementary Fig. S7**). Body weight was unrelated to active chases (**Fig. 4E**, *ρ =* 0.09, p = 0.235), but heavier animals were less likely to be chased (*ρ = -*0.34, p < 0.001).

In summary, social interactions in the NoSeMaze revealed distinct, quantifiable differences in chasing and competition-based social rank, which remained stable within a NoSeMaze round. Incidental tube competition outcomes established reliable social hierarchies, while chasing reflected individual differences in proactive social behavior.

### Stability of individual behaviors across social contexts

To assess whether behavioral features persist beyond a single NoSeMaze round, we examined their stability across rounds with different group members. More specifically, we quantified across-round stability with Spearman’s correlations between rounds 1 and 2 and with intraclass correlation coefficients (ICCs) computed across all rounds. High between-round correlations and high ICCs would indicate that these behaviors are dominated by stable between-mouse differences rather than transient adjustments to changing social contexts.

We found strong correlations between rounds for social rank and chasing. Specifically, stability was high for the tube competition-based David’s score (**Fig. 5A**, *ρ =* 0.57, p < 0.001), as well as for the fraction of active chases (**Fig. 5B**, *ρ =* 0.75, p < 0.001) and of times being chased (**Fig. 5C**, ρ = 0.54, p < 0.001). Notably, the fraction of active chases showed slightly higher across-round stability than tube-derived David’s score and the fraction of being chased. This is in line with active chases capturing an individual propensity to initiate this behavior, whereas competition-based social rank is a relational measure that is also influenced by the set of competitors and interaction opportunities in each reshuffled group. Nonetheless also social rank and being chased were overall stable. Across the full series, ICCs for these social measures were also in the good–excellent range, indicating high across-round stability (**Fig. 5D**, ICC = 0.548 to 0.890).

**Fig. 5:**
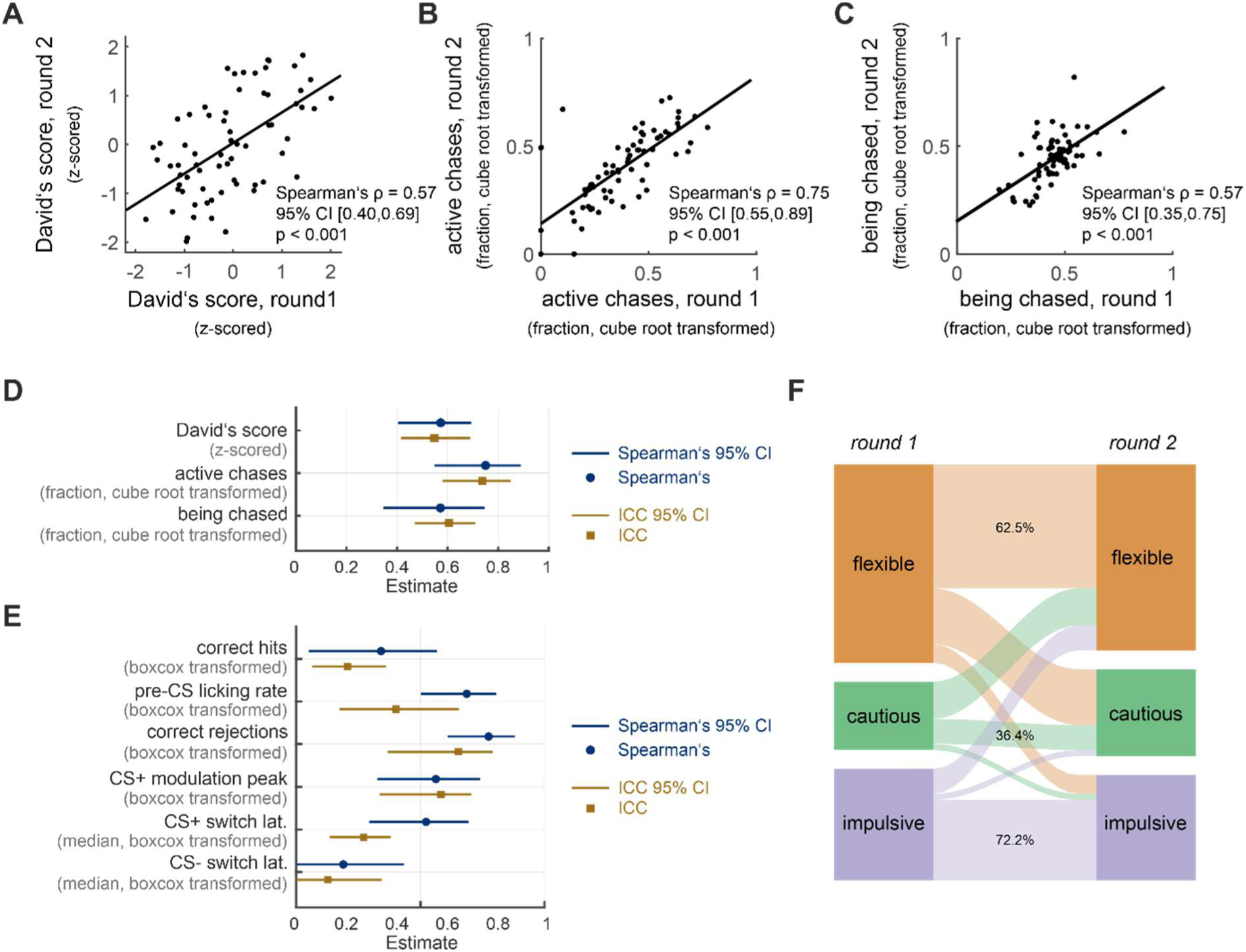
Stability of social rank, chasing, and reward-seeking features across repeated NoSeMaze rounds. **A–D**, Social rank and chasing were stable across NoSeMaze rounds. Individual David’s scores (**A**), fraction of active chases (**B**), and fraction of times being chased (**C**) were all significantly correlated between round 1 and 2 (Spearman’s ρ = 0.57, 0.75, and 0.54, respectively; all p < 0.001). Lines in A–C are illustrative only; inference is based on Spearman’s correlation. (**D**) Intraclass correlation coefficients (ICC) confirm similar stability for the three metrics across all rounds. Blue circles show Spearman’s ρ between rounds 1 and 2 with 95% CIs; golden squares show ICC with 95% CIs estimated across all rounds to quantify across-round stability of each metric. **E**, Stability of key reward-seeking features. As in **D**, blue circles depict Spearman’s ρ between rounds 1 and 2 (95% CIs), whereas orange squares give ICCs computed across all rounds (95% CIs) to index stability over the full longitudinal series. Features include correct hits and correct rejections, pre-CS licking rate, CS+ modulation peak, and switch latencies for CS+ and CS− (see also **Supplementary Fig. S8A-C**). **F**, Sankey diagram illustrates consistency in individual reward-seeking strategies across rounds. Most mice retained their behavioral strategy classification (flexible, cautious, or impulsive) between rounds, with the majority of flexible and impulsive animals remaining in the same category (see **Supplementary Fig. S8D**). CI, confidence interval; CS+, rewarded conditioned stimulus; CS−, unrewarded conditioned stimulus Note: Regression lines are illustrative only; statistical inference is based on Spearman’s correlation, which does not assume linearity.

For reinforcement learning, some features were stable across rounds while others were not (**Fig. 5E**). Correct hit rates correlated only moderately between rounds (ρ = 0.34, p = 0.008), with even lower ICCs (ICC = 0.21). In contrast, pre-CS licking (ρ = 0.69, p < 0.001, **Supplementary Fig. S8A**) and correct rejection rates (ρ = 0.78, p < 0.001, **Supplementary Fig. S8B**) showed higher between-round correlations but only moderate ICCs. Despite shifts in absolute levels, the pattern is consistent with their interpretation as stable individual styles related to impulsivity and inhibitory control (cf. **Fig. 5E**). Reward sensitivity, measured by peak CS+ licking modulation during anticipation, remained also stable across rounds (ρ = 0.56, p < 0.001, cf. **Fig. 5E**, **Supplementary Fig. S8C**) with a similar ICC. In contrast, the stability of CS+ and CS− switch latencies varied substantially. CS+ switch latencies exhibited low-to-moderate between-round correlations (ρ = 0.16 to 0.45 across reversals; median: ρ = 0.50) and only fair ICCs, whereas CS− switch latencies showed no reliable between-round correlation and poor ICCs (**Fig. 5E**). The generally shorter switch latencies in the second round (**Supplementary Table S3**) are consistent with meta-learning of the reversal rule, with a stronger effect on suppressing responses to CS− than on re-learning CS+. All statistical details are provided in the **Supplementary Material – Systematic Statistical Reporting**.

Cluster assignments across rounds showed moderate-to-strong contingencies (χ² = 25.51, p < 0.0001, Cramer’s V = 0.4573, **Fig. 5F**). Flexible learners, and especially impulsive go-learners, remained mostly stable across rounds (contingencies: 62.5% and 72.2%, respectively, **Supplementary Fig. S8D**). In contrast, cautious no-go learners were more likely to shift (contingency: 36.4%).

Together, convergent evidence from Spearman’s correlations between the first two rounds and ICCs across all rounds indicates that social rank, chasing, and reward-seeking features including impulsivity, inhibitory control, and reward sensitivity measured in the NoSeMaze represent stable individual styles.

Because the number of rounds in the NoSeMaze was unbalanced between animals (**Supplementary Table S3**) and age varied across groups (**Supplementary Table S1-2**), we also performed robustness checks. Stability estimates changed only minimally when recomputed in age-adjusted ICC models (between-group mean age and within-group age deviation; see Methods) and when restricting analyses to a balanced first-two-session subset (sessions 1–2 only; mice with both sessions) (**Supplementary Table S4**). Mixed models indicated that some chasing and reinforcement-learning measures showed modest age- and/or session-related shifts in absolute levels (**Supplementary Table S5**), but importantly, these did not affect the observed stability patterns.

As a secondary analysis, we tested whether OXTR^ΔAON^, which impairs de novo social recognition memory required for social clique formation in this cohort ^36^, also affects the measures reported here. Consistent with largely internalized features, OXTR^ΔAON^ produced only transient effects. Social rank was significantly lower during the first week in OXTR^ΔAON^ mice, but normalized over the following two weeks (**Supplementary Fig. S9A**), and no effects of OXTR^ΔAON^ were observed on chasing (**Supplementary Fig. S9B-C**) or reinforcement learning (**Supplementary Fig. S10**). In line with this, all reported stability relationships remained robust and statistically significant when controlling for OXTR^ΔAON^ (see **Extended Data Main**).

### Relationship between social rank, chasing, and reward-seeking features

Finally, we examined whether and how competition-based social rank, chasing, and cognitive styles were related, both at the individual and group level. We first compared social rank and chasing. Notably, David’s scores from tube competitions showed a moderately positive association with the fraction of active chases (**Fig. 6A**, ρ = 0.34, p < 0.001), suggesting that mice of high social rank were more likely to initiate chases. However, this association varied substantially across groups (range: ρ = –0.03 to ρ = 0.62, cf. **Fig. 6B**). We thus asked whether the strength of this relationship depended on how clearly the group’s social hierarchy was structured. Indeed, at the group level, the strength of the association between David’s scores and active chases was negatively related to transitivity (**Fig. 6B**, ρ = −0.64, p < 0.001), indicating that in groups with less clearly defined social hierarchies, active chases aligned more strongly with social rank from tube competitions. This suggests that aggressive signaling to assert social position becomes more important when dominance-subordination relationships are ambiguous. Conversely, in highly transitive groups, where social hierarchies were well established, active chases were less closely aligned with social rank, implying that social structure reduces the need for proactive reinforcement of dominance. Supporting this interpretation, the fraction of active chases performed by the mouse with the highest social rank also decreased with higher group transitivity (**Supplementary Fig. S11**, ρ = –0.61, p < 0.001).

**Fig. 6:**
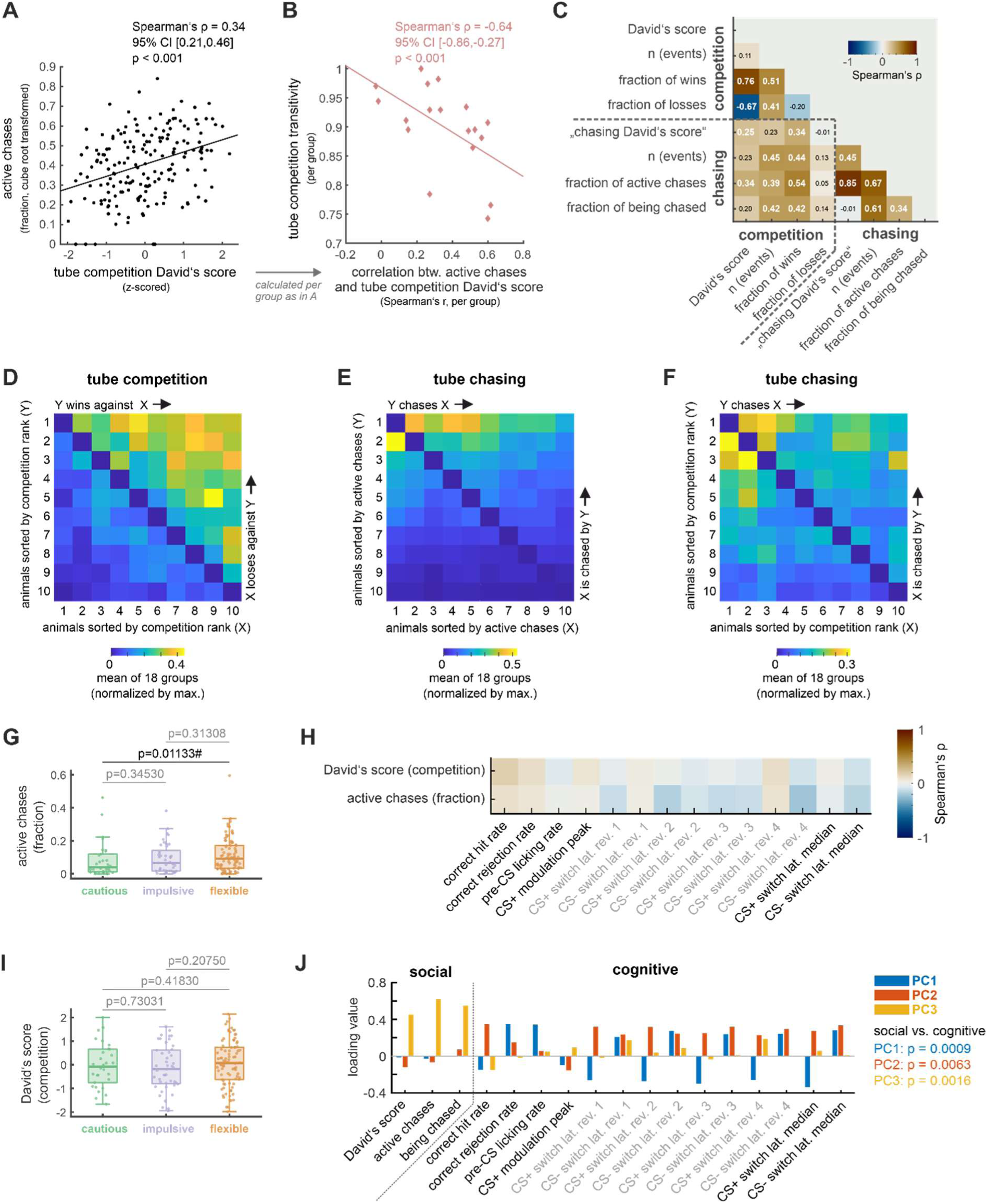
Relationship between social rank, chasing, and reward-seeking features. **A**, Social rank (David’s score) was positively correlated with the fraction of active chases. **B**, Across groups, the correlation coefficient calculated between David’s scores from tube competitions and fraction of active chases (x-axis, calculated separately within each group, as in **A**) was negatively associated with the group-level transitivity of the social hierarchy derived from tube competitions. Groups with lower transitivity showed a stronger association between social rank and chasing, suggesting that aggressive status signaling via chasing becomes more relevant when social hierarchies are less well-defined. **C**, Correlation matrix showing relationships between social rank and chasing metrics. Color scale indicates Spearman’s correlation coefficients; statistically significant values (Bonferroni-corrected) are shown in white. Note that the David’s score from tube competitions correlated to both, fraction of wins and losses in tube competitions, while the ‘chasing David’s score’ was strongly correlated only to the fraction of active chases, but not to the fraction of times being chased. **D-F,** Wins and losses in tube competitions (**D**) were concentrated in the top-right, occurring most consistently between animals of opposing social ranks. In contrast, chasing interactions (**E**, **F**) clustered in the top-left, and thus mostly within a subset of mice with high social rank (**F**), in which they may help clarifying dominance. **G**, Comparison of the fraction of active chases between the three different clusters defined from the reward-seeking task. Cautious mice initiated less chases than flexible individuals (p-values from permutation tests with n = 100,000 permutations using the median). Box plots show the median (line), interquartile range (IQR, box), and whiskers extending to 1.5 × the IQR. P-values were corrected for multiple comparisons using the Benjamini–Hochberg false discovery rate (FDR) procedure (α = 0.05). Significant comparisons after FDR correction are marked with #. **H**, Correlation heatmap between David’s score (social rank from tube competitions), active chases, and reward-seeking features showed no significant associations after FDR correction for multiple comparison. **I**, Comparison of the David’s score (social rank from tube competitions) between the three different clusters defined from the reward-seeking task showed no significant difference (p-values from permutation tests with n = 100,000 permutations using the median). Box plots show the median (line), interquartile range (IQR, box), and whiskers extending to 1.5 × the IQR. **J**, Loading values from a principal component analysis of social rank (David’s score), chasing (active chases and being chased), and reward-seeking (cognitive) features for the top three components (PC1–PC3). Bars indicate each feature’s contribution to the respective component. PC1 and PC2 were dominated by cognitive features (p = 0.0009 and p = 0.0063), PC3 by social rank and chasing (p = 0.0016; permutation tests, n = 10,000). The largely non-overlapping loadings suggest orthogonality between social and cognitive domains. CI, confidence interval; CS+, rewarded conditioned stimulus; CS−, unrewarded conditioned stimulus Note: Regression lines are illustrative only; statistical inference is based on Spearman’s correlation, which does not assume linearity.

To gain further insight into the interplay between social rank and chasing, we examined the structural properties of both behaviors in more detail. When considering only social rank, the David’s score correlated strongly with both the fraction of wins (ρ = 0.76, p < 0.001) and the fraction of losses (ρ = −0.67, p < 0.001) in tube competitions (**Fig. 6C, Supplementary Fig. S12A-B**). The integrated tube tests thus meet the assumption of symmetry, where both dominant pushing and subordinate retreat equally contribute to social rank ^1,37^.

Tube crossings in the NoSeMaze are motivated by the intent to eat, drink, sleep, or socialize. Accordingly, competitions within the tube arise by chance when two animals enter from opposite sides at the same time. Social rank therefore captures the consistent outcomes of repeated incidental competitions (push versus retreat). This measure is not confounded by differences in tube engagement, as social rank was neither associated with participation in tube competitions (ρ = 0.11, p = 0.139; **Fig. 6C, Supplementary Fig. S12C**) nor with the overall number of tube detections when controlling for chasing (Spearman’s partial correlation between detection count and z-scored David’s score, corrected for the fraction of active chases: ρ = 0.022, p = 0.758).

In contrast, these assumptions were not met for chasing (**Fig. 6C**). This is best illustrated by constructing a ‘chasing David’s score’. This score was positively associated with the total number of chasing events (ρ = 0.45, p < 0.001, **Fig. 6C, Supplementary Fig. S12D)**, reinforcing that chasing reflects volitional, actively initiated behavior. Importantly, the constructed ‘chasing David’s score’ also correlated strongly with the fraction of active chases (ρ = 0.85, p < 0.001, **Fig. 6C, Supplementary Fig. S12E**) but was entirely unrelated to the fraction of times being chased (ρ = −0.01, p = 0.897, **Fig. 6C, Supplementary Fig. S12F**). This fundamental difference suggests that, within a group, chasing did not follow a symmetric dominance-subordination gradient as seen in social rank. Instead, it reflected an unbalanced distribution of initiators and recipients. Consistently, the relationship between the fractions of active chases and being chased differed significantly from the relationship between the fraction of wins and losses in the tube competition (cf. **Supplementary Fig. S6A-B, Supplementary Fig. S13**). As a result, the fraction of active chases provided a more direct measure of volitional engagement in chasing than its David’s score, as it directly quantified proactive initiation.

With this distinction in mind, we further explored the specific relationship between incidental tube competitions and active chases (**Fig. 6D-F, Supplementary Fig. S14**). The distribution of wins and losses in incidental tube competitions (**Fig. 6D, Supplementary Fig. S14A**) differed from those of active chases and being chased (**Fig. 6E, Supplementary Fig. S14B**). Specifically, a subset of mice expressed high levels of both active chases and being chased (**Fig. 6F**), and this subset preferentially mapped onto mice with high social ranks in tube competitions (top left cluster, **Fig. 6F, Supplementary Fig. S14C**). These findings suggest that chasing may serve to negotiate relative social rank among animals with high social rank.

Finally, we examined how social rank and chasing mapped onto reward-seeking strategies. Cautious no-go learners were less likely to engage in active chases compared to flexible learners (**Fig. 6G**, permutation test with 10,000 permutations, p = 0.01133). Apart from this, no significant group differences were found for competition-based David’s score (cf. **Fig. 6I**, see **Supplementary Material – Systematic Statistical Reporting** for details) or fraction of times being chased (**Supplementary Fig. S15**). Moreover, neither social rank nor active chases were associated with individual reward-seeking features, including impulsive pre-CS licking, reward sensitivity, inhibitory control, or cognitive flexibility (**Fig. 6H**, for Spearman’s ρ, raw-and corrected-p-values, see **Supplementary Material – Systematic Statistical Reporting**). Consistent with this dissociation, principal component analysis showed that social and cognitive domains loaded onto distinct components, further supporting their independence (**Fig. 6J**, permutation test (10,000 permutations) on the mean loadings values; PC1: p = 0.0009, PC2: p = 0.0063, PC3: p = 0.0016). Notably, all reported between-metric relationships remained robust and statistically significant when controlling for OXTR^ΔAON^ (see **Extended Data Main**).

Together, these findings suggest that the here examined non-social cognitive features are largely orthogonal to the individuals’ social rank and propensity to chase.

## DISCUSSION

Ecologically enriched, yet experimentally controlled assessments allow us to study behavioral individuality and social structure in group-living animals over extended timescales. Here, we show that individual mice carry stable, individual-specific, and multi-faceted profiles of social position and cognitive styles across changing group contexts. Our approach goes beyond prior work by testing cross-context stability under repeated, systematic group reshuffling in larger mouse societies, while measuring competition outcomes, chasing, and reinforcement-learning behavior in parallel. To capture this, we had developed the NoSeMaze, an open-source modular platform for continuous tracking of spontaneous behavior in socially housed mice. Although ecological approaches are increasingly valued in neuroscience ^17^, few existing systems support long-term, automated monitoring of individuals across multiple behavioral domains in larger mouse groups. Most current paradigms are tailored to specific behaviors, such as aggression ^24,25^, conflict ^38^, or locomotor activity ^30^, and offer only limited integrative capacities ^12,19,39–41^. The NoSeMaze addresses this gap by the simultaneous, longitudinal assessment of social rank, chasing, and olfactory reinforcement learning. Its modular, open design supports future expansion to other behavioral domains and integration with neurobiological tools, providing a flexible framework for studying social structure and behavioral individuality. This study focused primarily on the relation of social rank and chasing. We however also considered their relation to additional variables including the loss of oxytocin receptors in the olfactory cortex in the adult (OXTR^ΔAON^), involved in de novo social recognition learning. The propensity to chase was largely unaffected by OXTR^ΔAON^. Mice carrying OXTR^ΔAON^ displayed a transient reduction in social rank during the first week that normalized thereafter. This transient effect contrast to the persistent impairment by OXTR^ΔAON^ in forming higher-order social bonds that enable membership in stable cliques, as identified by video tracking of self-paced interactions in the same cohort ^36^. Together, these findings suggest that OXT-dependent olfactory learning is critical for the formation of social context-dependent higher-order bonds, but plays a limited role in shaping hierarchy-related behaviors.

Social hierarchies ^1,42^ were assessed from incidental dyadic interactions in the novel integrated tube-test module. This method enabled social rank estimation without experimenter intervention and was validated against manual tube testing. Importantly, participation frequency and differences in tube entry time did not confound the resulting social ranks, underscoring the robustness of the automated incidental rank assessment. Social rank was equally shaped by dominance and subordination in tube competitions. This supports Bernstein’s assertion ^37^ that social ranks are maintained not only by dominance but also by voluntary subordination. While this study focused on male mice, in which social hierarchies are best established ^9^, future work is needed to explore sex-specific expressions of social structure and their neurobiological underpinnings in female groups. We therefore restrict our interpretation to males. Critically, male social ranks were robustly maintained within the same group over time. Even more interestingly, the longitudinal accelerated design, in which animals lived in different groups composed of different individuals, allowed us to test whether social rank is predominantly driven by current group composition or reflects an internalized component that generalizes across social contexts. Indeed, individuals occupied similar relative social ranks across time and groups with different members. This temporal and contextual stability supports interpreting social rank as a stable, internalized characteristic of individuals. In this sense, ‘internalized’ refers to stability across repeated rounds and reshuffled groups in this paradigm and to relatively stable tube-competition outcomes. The tube-derived social rank describes here a dimension of individual social behavior. The capacity to quantify stable individual differences across changing social contexts highlights the value of the NoSeMaze for lifespan-oriented studies of behavioral individuality. A future direction is to extend this framework to earlier developmental stages to understand which early experiences shape later trajectories of social position.

In more constrained or despotic conditions, chasing can serve as a unidirectional, dominance-related behavior directed at subordinates ^9,43,44^. The larger groups observed in the NoSeMaze reveal a more nuanced role for chasing behavior. Chasing levels are individually stable, but the expression and meaning of chasing are context-sensitive. Chasing was neither broadly distributed nor consistently directed down the social hierarchy. Instead, it was initiated by a small subset of individuals – primarily those occupying high competition-based social ranks – and frequently occurred reciprocally within this group, suggesting intra-elite social dynamics rather than broad dominance enforcement. Rather than solely serving to impose social hierarchy, chasing appeared to function as a means through which individuals with high social rank monitor, negotiate, and maintain their relative standing within the top tier. Notably, the identity of frequent chasers remained stable over time and persisted across changing group compositions, indicating that the propensity to initiate chases reflects a consistent individual-level tendency rather than a purely situational response.

However, its coupling to social rank depended on the group’s hierarchy structure. In groups with less clearly defined social hierarchies (i.e., lower transitivity), active chases aligned more strongly with social rank, suggesting that mice in less structured groups rely more on proactive signaling to clarify social rank. Indeed, the top-ranked mice in these groups exhibited relatively high levels of active chases. This context-sensitivity highlights chasing’s dual role: it serves as a tool for negotiating social rank among mice at the upper end of the hierarchy, and additionally functions to establish or reinforce hierarchical clarity when social structures are ambiguous. These dynamic aspects of chasing, including its asymmetric initiator–recipient structure and proactive engagement, differ from the nature of tube competitions, which are incidental encounters. Together, tube-derived social rank and chasing describe complementary dimensions of social position, and, alongside other features such as clique formation ^36^, contribute to describe facets of a broader multidimensional social behavior. Within this complex environment, chasing emerges as a flexible behavioral propensity that is dissociable from formal social rank. Specifically, in the NoSeMaze, chasing contributes dynamically to the maintenance, negotiation, or clarification of social hierarchy structure. These findings reveal a novel aspect of social dynamics: chasing is not merely a dominance display but a flexible context-dependent mechanism shaped by both individual disposition and group-level social structure.

In addition to behavioral strategies, physical features can also influence patterns of social interaction. Body weight is a factor implicated in social rank and aggression ^45^, yet studies in mice have yielded mixed results ^2,46^. In the NoSeMaze, heavier animals were less likely to lose tube competitions or to be targeted in chases, but they were not more likely to win tube competitions or initiate chases. This suggests that weight may buffer against subordination rather than confer dominance, acting as a passive protective factor that reduces the likelihood of being challenged by others.

Beyond social and physical characteristics, we also assessed reward-guided cognitive features, including impulsivity, reward sensitivity, and inhibitory control. These features were relatively stable considering the dynamics of reinforcement learning. Mice adopted distinct cognitive styles: impulsive ‘go’, cautious ‘no-go’, and flexible learners. The styles that were most effective in retrieving rewards also showed the highest stability over months. The cognitive profiles also map onto known axes of behavioral variation observed across species ^47–50^. Although social and cognitive domains revealed internally consistent profiles, they were in many cases orthogonal to each other. This is in line with meta-analyses showing weak associations between social position and cognitive characteristics in mice ^14^. However, when viewed as composite styles, some links emerged: cautious ‘no-go’ learners were least likely to chase others, while flexible learners showed the highest levels of proactive chasing. Contrary to expectation, impulsivity did not reliably predict active chases, suggesting that social engagement in dominance negotiation may require flexible rather than impulsive behavior.

Together, these findings highlight the value of the NoSeMaze as an integrative ecological platform for dissecting social group organization and behavioral individuality. Different dimensions of social position map onto well-established factors in resilience. The approach may thus provide a suitable testbed for modeling and causally dissecting resilience mechanisms. The NoSeMaze aims to increase environmental complexity and group dynamics while retaining experimental control, enabling longitudinal high-dimensional phenotyping of individuals. At the same time, it is a controlled laboratory group-housing habitat optimized to capture a subset of the determinants of social complexity present in natural communities. The strength of this design lies in the continuous, observer-independent observation of complex behaviors in defined environmental and social contexts. Accordingly, we interpret our findings as applying to the social contexts and environmental conditions tested here. Building on this, its modular design allows for introducing additional environmental and social factors like stressors, mating behavior, or resource competition in the future to understand their respective impact in modifying social behaviors. Further, the NoSeMaze is compatible with longitudinal, individualized neuroscience, including chemogenetic ^51^ or pharmacological circuit manipulations that can be applied in a time-locked, subject-specific manner via the lickports. The NoSeMaze can also be combined with intermittent task-based fMRI protocols, mirroring study designs currently emerging in human neuroscience ^52,53^. Together, this makes the NoSeMaze suited for unraveling the neural, genetic, and environmental contributions to individuality over time and across contexts.

## Conclusion

The NoSeMaze provides a novel and powerful platform for studying the emergence of social structure and behavioral individuality in mouse societies. By enabling continuous, automated tracking of social hierarchy, chasing, and cognitive strategies within the same ecological context, the NoSeMaze revealed that individual behavioral features are remarkably stable over time, even across changing group compositions. This suggests that social position is not merely a product of immediate social environments, but rather reflects a stable individual characteristic that persists across different social contexts.

Crucially, social position is not fully described by a single behavioral dimension. We focused here on two separable dimensions: competition-based social rank and proactive chasing. Future work should integrate additional dimensions of social organization, such as affiliative bonding and higher-order network measures ^36^, to capture additional aspects of social organization. Chasing behavior played a dual role, reflecting a stable individual propensity while also adapting to group-level structure. It was most pronounced among high-ranking individuals and aligned more closely with social rank when hierarchies were less well clarified, suggesting that chasing serves both stable and context-sensitive functions. At the same time, social and cognitive domains were largely independent, challenging the idea of unified behavioral types and supporting a multidimensional view of individuality.

Finally, the NoSeMaze exemplifies how the complexity of group behavior in semi-naturalistic conditions can be captured without sacrificing experimental control. Its modular design and compatibility with longitudinal neural interventions provides a tool for linking behavioral characteristics to neural circuit dynamics, genetic factors, and environmental influences. By combining high-dimensional phenotyping with ecological validity, the NoSeMaze opens new avenues for understanding how stable individual differences in social and cognitive function arise and how they shape, and are shaped by, the brain across time and context.

## METHODS

### Animals

#### Animal strain and housing conditions

A total of 79 adult male homozygous OXTRfl/fl mice (B6.129(SJL)-*Oxtr^tm1.1Wsy^*/J, RRID: IMSR_JAX:008471, Jackson Laboratory) backcrossed > F10 to C57BL/6J background (Charles River, Sulzfeld) were used for the experiments. Animals entered the NoSeMaze as adults (see **Supplementary Table S2** for ages at NoSeMaze entry across rounds) and had been continuously group-housed after weaning (3–5 mice/cage), i.e., they were not developmentally or socially naïve at study onset. Of the 79 mice, 26 were injected six weeks before the start of the experiment with an AAV expressing Cre recombinase (*rAAV1/2-CBA-Cre*) into the AON pars centralis to induce bilateral OXTR deletion (OXTR^ΔAON^). The remaining 53 animals received an AAV expressing only dTomato (*rAAV1/2-CBA-dTomato*). Details on viral constructs, stereotactic surgery, and injection procedures are provided in Wolf et al. ^32^. Genotype-dependent effects are mainly addressed in a separate manuscript ^36^. Each mouse was implanted in the neck with a subcutaneous RFID chip (Euro I.D. ID 100 trovan®) for individual identification using a microchip syringe injector under brief anesthesia with 1.5% isoflurane. During home-cage housing, mice were kept in groups of 3-5 under a 12h light/dark cycle (room temperature 24 °C, air humidity 55%). Food and water were provided ad libitum. Before first entry into the NoSeMaze, home-cage group composition was reshuffled every 3–4 days over several weeks to ensure familiarity among potential future group members (cf. **Fig. 1C-D**). During the experiment, mice were tested across multiple NoSeMaze rounds and were systematically reassigned to different NoSeMaze groups between rounds (cf. **Fig. 1D**), while remaining within their respective age cohort. This repeated reshuffling of NoSeMaze group composition was a central feature of the study and enabled us to test whether individual differences in social rank, chasing, and reinforcement-learning behavior generalized across changing social contexts. During the intervals between NoSeMaze rounds, mice were also reassigned to different home-cage groups. Importantly, both home-cage group assignments and NoSeMaze group compositions were randomized and blinded to prior behavioral outcomes or individual performance. In the NoSeMaze, animals had free access to food. Water was obtained from the self-paced stimulus-outcome learning task that was accessible 24/7. The number of trials determined the water intake, which was continuously monitored by the number of licks in go trials.

#### Ethics statement

All procedures were in accordance with the National Institutes of Health Guide for the Care and Use of Laboratory Animals and the EU 2010/63 directive, and approved by the local animal welfare authority (Referat 35, Regierungspräsidium Karlsruhe, Karlsruhe, Germany).

#### Sex

In this study, we used male mice, for which social hierarchy is best studied ^9^ and for which the NoSeMaze was originally designed and optimized. The cohabitation of male and female mice in the same NoSeMaze leads to mating and thus different dynamics in the social behavior. Future studies will leverage current efforts to determine optimal NoSeMaze conditions for female mouse societies to generalize the observations across sexes.

### Non-invasive sensor-rich maze (NoSeMaze) and behavioral data acquisition

#### NoSeMaze infrastructure and habituation

The NoSeMaze contains an olfactory learning module (**Fig. 1A** and **Supplementary Fig. S1**), which is controlled by an open-source Python software that we modified based on Erskine et al. ^40^. Full documentation is available at https://github.com/KelschLAB/NoSeMaze. Four identical NoSeMazes, each housing n=9-10 male mice, were used in parallel (for group sizes, see **Supplementary Table S2**). All mice were habituated to the NoSeMaze twice in subgroups of 6-7 animals. Each acclimatization period lasted 16-20 hours including one dark cycle, giving the animals enough time to explore the novel environment. The age ranges at which mice entered the NoSeMaze for the different rounds are shown in **Supplementary Table S2**.

#### Stimulus-outcome learning task at the water lick port

The task at the olfactory stimulus-outcome learning module, where the animals earned their daily water, was gradually introduced during acclimatization. Decanal (Sigma Aldrich W236217) and octanal (Sigma Aldrich W279706), both diluted in mineral oil, were used as odors and applied at a final concentration of 0.1 % with a custom-built olfactometer. Initially, a drop of water (10 µl) was given each time the lick port was visited. Then, only one of the two odors was associated with a water drop. The odor presentation started 500 ms after the animal was recognized by its RFID chip and lasted for 2 s (cf. **Fig. 2A**). When the mice continuously lived in the NoSeMaze, in the go trials, one odor was followed by a water drop in 100% of the trials if the animal licked twice during the odor presentation. In the no-go trials, the other odor (CS−) was presented and not rewarded. If the animals licked more than once during odor presentation, they received a timeout of 6 s to initiate the next trial. The reward contingencies switched every three days (cf. **Fig. 2B**). The number of trials and licks were registered with a Darlington sensor and assigned to the RFID tag of each mouse recognized by the antenna at the lick port. Together with the licking behavior, the water intake per animal was recorded and controlled daily. All animals consumed at least 2 ml (up to 6 ml) water per day.

#### Tube competitions and chasing

Two tubes connected the housing area and the social arena, where the access to the water port was located. Thus, mice had to cross the tubes regularly to consume both food and water. Custom written MATLAB scripts detected events of tube competitions and tube chasing. A competition event was defined as a sequence of RFID-tag detections indicating that two mice entered the tube from different sides at the same time and one mouse pushed the other back out of the tube. While the ‘dominant’ mouse was detected on both sides of the tube, the ‘subordinate’ mouse was detected twice at its entry detector (when entering and when exiting backwards, see **Fig. 3A**). Detection patterns of putative competition events were visually inspected and included in the analysis if they met the criteria outlined above. A ‘chasing event’ was defined as a sequence of detections indicating the simultaneous presence of two mice in the tube, moving in the same direction and in close succession (**Fig. 4A**). Events were classified as chases if (1) the time difference between the two animals at the detector was less than 1.5 seconds (Δtime at detector < 1.5 s), and (2) both animals crossed the tube within 2 seconds (Δtime through tube < 2 s). Animals were visually inspected daily for their well-being. Notably, we did not observe any intensive fighting or relevant injuries during the entire duration of the experiment in any of the groups.

### Quantification of individual reward-seeking behavioral metrics

#### Metrics derived from the stimulus-outcome learning task

A detailed summary of the data across NoSeMaze rounds is provided in **Supplementary Table S1**. Data from 17 NoSeMaze groups were used for the analysis of the reward-seeking domains (see **Supplementary Table S2**). In four NoSeMaze groups, the paradigm at the reinforcement learning port did not work properly, so that they had to be excluded from the analyses. Six animals, distributed across different groups, were additionally excluded due to insufficient RFID recognition at the lickport. The variability in licking responses was quantified using the following metrics: (1) correct hit rate, (2) correct rejection rate, (3) pre-CS licking rate before odor presentation (from +0.05 to +0.5 s relative to trial start; see **Fig. 2G**), (4) peak licking during the CS+ window (from +1 s to +2.4 s relative to trial start; ‘CS+ modulation peak’, **Fig. 2G**), (5) switch latencies for CS+ and CS−, computed separately for each reversal and summarized as the median across reversals, and (6) the number of trials required for animals to achieve an 80% correct hit or correct rejection rate (‘time to criterion’). All trials were used to calculate pre-CS licking rate, correct hit rate, and correct rejection rate. The pre-CS licking rate served as a measure of impulsivity, while licking inhibition during CS− trials reflected inhibitory control. Reward sensitivity was assessed using CS+ modulation peak, defined as the maximum licking rate during CS+ presentation, by concatenating the last 150 CS+ trials before each reversal to ensure that animals had reached a stable state. CS+ switch latency quantified how quickly an animal increased licking behavior in response to the newly rewarded conditioned stimulus (CS+) after a reversal. It was defined as the number of CS+ trials needed after a reversal for the animal to return to at least 70% of its pre-reversal CS+ licking rate. A shorter CS+ latency reflected faster acquisition of the new reward contingency. CS− switch latency captured how quickly an animal suppressed licking in response to the newly unrewarded conditioned stimulus (CS−) after a reversal. It was calculated as the number of CS− trials after a reversal needed for licking to drop below 50% of the pre-reversal CS+ licking rate. A shorter CS− latency indicated faster response inhibition or suppression of previously reinforced behavior. For both metrics, the pre-reversal licking rates were computed from the last 150 CS+ trials preceding each reversal. All rates were calculated from the odor presentation window (0.5-2.5 s). If no switch was detected, the switch latency was set to the total number of available post-reversal CS+ or CS− trials, which varied by animal and phase. Note that both metrics, CS+ and CS− switch latencies, reflect individualized measures, as they referred to each mouse’s own pre-reversal licking rates. Lastly, time to criterion was assessed as the number of trials following a reversal required for the animal to achieve at least 80% correct responses – hits for CS+ and correct rejections for CS− – sustained over a 10-trial moving average. If the animal did not reach this performance level during the post-reversal phase, the time to criterion was set to the total number of available post-reversal trials for the respective stimulus, indicating that the performance criterion was not met within the phase. In contrast to the switch latencies, time to criterion referred to a uniform performance threshold for all animals. For illustrative purposes, some of the data were cube root or Box-Cox transformed in cases of non-normal distribution.

#### Clustering of reward-seeking profiles

Data from six features were used for the clustering approach: CS+ and CS− median switch latencies, correct hit rates, correct rejection rates, CS+ modulation peak, and pre-CS licking rates. All features were standardized using min-max normalization to preserve individual ranges despite potential skew. Then, K-means clustering was applied. The optimal number of clusters (k = 3) was determined using the elbow method. A silhouette score of 0.50 indicated a moderately good separation between clusters. Cluster assignments were visualized in PCA-reduced space using the first three principal components, and statistically compared across features using permutation tests. To assess differences of the six reward-seeking features between clusters, we performed unpaired permutation tests using 100,000 iterations and a median-based test statistic to account for data skewness. To address repeated measures and within-animal dependencies, a realistic swap ratio was estimated by computing the ratio of between-animal to total variability (i.e., between-animal / [within-animal + between-animal] variability). This swap ratio defined the proportion of data globally shuffled across clusters during each permutation. It was calculated per feature within each cluster and was applied globally during permutation tests to maintain realistic within-subject structure. Specifically, we used the mean of the two swap ratios from the two clusters that were compared. By incorporating both within- and between-subject variability, this approach generated a more robust null distribution. Final p-values were computed using two-tailed tests comparing observed and permuted differences. To control for multiple testing, p-values were adjusted using the Benjamini–Hochberg false discovery rate (FDR) procedure ^54^.

### Quantification of individual social behavioral metrics

#### Social rank and chasing metrics

Individual-level social rank was quantified using the David’s score from tube competitions. It was calculated based on the counts of wins and losses between all pairs of animals. Specifically, for each pair of animals *i* and *j*, we calculated the number of times that *i* defeated *j* (*a*_*ij*_) and the total number of encounters between them (*n*_*ij*_ = *a*_*ij*_ + *a*_*ij*_). The directed proportion of wins was then defined as *P*_*ij*_ = *a*_*ij*_/*n*_*ij*_ whenever *n*_*ij*_ > 0 For each individual *i*, the first-order scores were calculated as

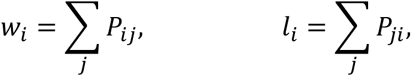

representing the summed proportions of wins and losses, respectively. Pairs with no interactions (*n*_*ij*_ = 0) produce undefined *P*_*ij*_ values, which were excluded from the sums. To capture the strength of opponents, we additionally computed second-order terms, where

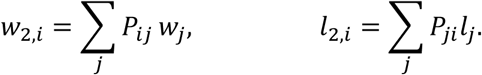

These weighted proportions assign greater value to a win against a high-ranking opponent than against a low-ranking one, and analogously for losses ^55^. The David’s score for each individual was then obtained as

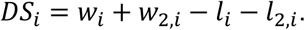

Linear ranking of David’s scores within a group yielded each animal’s social rank, with 1 denoting the highest social rank. In this study, we determined the social rank for every animal based on the first three weeks in the NoSeMaze. Similarly, a ‘chasing David’s score’ was calculated using the counts of active chases and times being chased between all pairs of animals in place of wins and losses. Moreover, we computed the fraction of active chases and the fraction of times being chased for each animal, defined as the number of actively initiated chases or times being chased divided by the total number of chasing events in its group. Similarly, the fractions of wins and losses in tube competitions were calculated for each individual based on the outcomes of these events.

#### Group-level characterization of social hierarchy structure

Social hierarchies were evaluated using metrics recommended in recent literature ^35^, including steepness, transitivity, stability, and uncertainty-by-repeatability (**Fig. 3C**, see also ‘Source Data’). Triangle transitivity was used to quantify transitivity of the social hierarchy by calculating the proportion of transitive triangles (i.e., triads of animals) from all triangles ^56^. A triangle is transitive when, if A dominates B and B dominates C, then A also dominates C. The triangle transitivity index therefore reflects the fraction of all triads in the group that meet this criterion. Significance of triangle transitivity was tested with a permutation test by computing 1,000 random networks (while preserving the distributions of observed null, tied and asymmetric edges in the hierarchy network). These random networks’ triangle transitivity values were compared against the observed values ^56^. Steepness describes the degree of dominance success over subordinate animals and is defined as the slope of a linear fit of normalized David’s scores ^57^. The significance of a hierarchy’s steepness was also assessed with a permutation test, in which 1,000 hierarchies with wins and losses assigned by chance are calculated and compared to the observed steepness. The stability index ranges between 0 and 1, with 0 indicating unstable hierarchies, in which the ordering reverses every other day, and 1 describing stable hierarchies without rank changes ^35^. The uncertainty-by-repeatability is calculated as the intraclass correlation coefficients for each individual after randomizing the order of the interactions. A repeatability score above 0.8 suggests a reasonably robust hierarchy ^35^. The metrics were calculated in R using the packages ‘aniDom’ (https://cran.r-project.org/package=aniDom) ^35^ and ‘EloRating’ (https://cran.r-project.org/web/packages/EloRating/index.html) ^58^.

#### Validation of social rank metrics

To validate the integrated social rank metrics and demonstrate their convergent validity in assessing social rank, David’s scores derived from the integrated tube test were compared to two established dynamic social hierarchy estimation methods: the traditional Elo-rating and the randomized Elo-rating ^35^. Further, to cross-validate the integrated tube test readout, a conventional tube test was performed the day after mice were removed from the NoSeMaze and had returned to standard cages. The standard tube test was performed on ten NoSeMaze groups. In this standard setup, mice were habituated to a classical tube test apparatus consisting of two compartments connected by a tube with a central barrier. Random dyads were drawn from each group and a tube test was conducted by placing the two mice in the tube facing each other. If one mouse pushed the other out of the tube within one minute it was scored a win or loss respectively, otherwise a tie. Each animal performed five such trials to avoid exhaustion ^21^. The resulting David’s score from the manual tube tests were correlated significantly to those from the integrated tube test (**Supplementary Fig. S4C**). To assess the influence of entry-time differences in tube competitions, we computed two versions of dynamic Elo ratings for each animal based on its observed win/loss outcomes. The ‘uncorrected’ Elo rating was calculated using standard update rules, where each win/loss updated an individual’s Elo score based on the pre-match Elo ratings and a group-specific K-factor ^58^. To evaluate and correct for potential biases due to asymmetries in tube entry time, we first quantified the influence of entry-time differences on tube competition outcomes using a linear mixed-effects model (LME). For this, we compiled a dataset combining tube competition events across all groups. Each event was included twice – once from the perspective of the winner and once from the loser. Each row contained the individual’s and the opponent’s identity, the pre-match Elo ratings of both animals, and their entry times. From these, the differences in both Elo rating and entry time were computed and included as predictors. Extreme values for entry-time difference were capped at a threshold of 10 seconds to minimize the influence of outliers. The LME was implemented in R (v4.4.1) using the lme4 package ^59^ (glmer() function, binomial family). The binary outcome variable indicated wins or losses, with scaled Elo difference and scaled entry-time difference as fixed effects, and mouse identity as a random intercept. The estimated fixed-effect coefficient for entry-time difference was then used to adjust the expected win probability in the Elo update formula. Specifically, for each match, the scaled entry-time difference was multiplied by the LME-derived coefficient and added to the log-odds of the standard Elo expectation. This resulted in a ‘corrected’ Elo rating, which was updated step-wise throughout the match sequences in each group. Finally, corrected scores were rescaled to fit the distributional properties (mean and SD) of the ‘uncorrected’ Elo scores to ensure comparability across methods.

#### Statistics analysis

Note that full details on the statistical tests, including *n*, degrees of freedom, exact p-values and 95% confidence intervals, are provided in the **Supplementary Material – Systematic Statistical Reporting**, as well as in the **Extended Data** for the LMEs accounting for different covariates.

#### Basic data assessments

We quantified the number of trials at the lickport, the number of tube competitions, and the number of chasing events per group and day, and illustrated those using boxplots. The time of day at which licking and social interaction events occurred was visualized with polar plots.

#### Correlation and stability analyses of behavioral metrics

We used Spearman’s rank correlation coefficients (ρ) to examine relationships among behavioral metrics, as well as between behavioral metrics and body weight. To account for repeated measures, we also fitted LMEs with mouse identity as a random intercept (see **Extended Data Main** and **Extended Data Supplement**). Note that the LMEs confirmed all results from the correlation analyses shown in the manuscript. To assess the relationship between social rank, chasing, and body weight, the z-scored body weight (within groups) before entering the NoSeMaze was correlated with the David’s score from tube competitions, the fraction of active chases, and the fraction of times being chased. To evaluate the temporal stability of the social metrics within a NoSeMaze round, social metrics were calculated separately for the first, second, and third week, and cross-correlated across weeks. Most mice (n = 68) participated in at least two NoSeMaze rounds with reshuffled group members. **Supplementary Table S2** summarizes the number of rounds per mouse and missing data due to technical problems. We examined the stability of social and reward-seeking metrics by correlating values from the first and second round (Spearman’s correlation, cf. **Fig. 5**). These round-1-to-round-2 correlations use one paired observation per mouse and are therefore not inflated by mice contributing >2 rounds. Additionally, we assessed the consistency of cluster assignments based on reinforcement learning metrics between rounds using a chi-square test and visualized transitions with a Sankey diagram. Importantly, to quantify stability across all available rounds, we also estimated intraclass correlation coefficients (ICCs) from variance-component LMEs, interpreting ICCs as the proportion of total variance attributable to stable between-mouse differences. The ICC treated mouse identity as a random intercept and NoSeMaze group (i.e., round-specific social group) as a random effect, with repetition included as fixed effect (i.e., stability across groups while holding repetition means constant). Formally, for a given metric

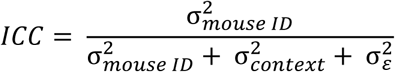

with σ^2^_context_containing the group as the relevant context term. The time between rounds extended over several weeks (mean: 6.8 weeks, SD: 2.8 weeks; see **Supplementary Table S2** for details). For correlation analyses, we obtained 95% confidence intervals by subject-level bootstrap (resampling mice with replacement; B = 2,000 draws; percentile intervals), preserving the within-mouse pairing of rounds. For ICCs, we computed 95% confidence intervals by subject-level bootstrap using the same resampling scheme and refitting the LME for each bootstrap sample.

Variance components for ICC estimation were obtained from LMEs fit by restricted maximum likelihood (REML), which yields less biased variance-component estimates and is well suited for unbalanced repeated-measures designs (i.e., different numbers of rounds per mouse). To assess potential confounding by age structure, age was decomposed into a between-group component (group-mean age) and a within-group component (each animal’s deviation from its group mean) and included as covariates. ICCs were recomputed in age-adjusted models (see **Supplementary Table S4**). Finally, we performed a conservative sensitivity analysis restricted to the first two sessions per mouse and to mice with observations in both sessions (“balanced first2”), to additionally account for unbalanced participation structure.

#### Comparative analysis of chasing and social rank

Distributions of the fraction of active chases and the fraction of times being chased, pooled across all groups, were assessed using histograms and empirical cumulative distribution plots (**Supplementary Fig. S6A**). Distribution shapes were compared via Kolmogorov–Smirnov tests. A similar analysis was applied to the distributions of wins and losses in tube competitions (**Supplementary Fig. S6B**). To further examine differences between the fractions of active chases and of times being chased, we computed cumulative sums over the sorted data within each group, visualized them using shaded error bar plots, and tested group-level differences between the two metrics using unpaired permutation tests (cf. **Fig. 4C**, 10,000 permutations). Lastly, to explore whether win-loss relationships differed between chasing and tube competition events, we combined data from both modalities and modeled the fraction of ‘wins’ – defined as successful tube competitions or active chases – using a LME (**Supplementary Fig. S13**). Active chases were treated as ‘wins’ in the context of chasing, being chased as ‘losses’. The model included the fraction of ‘losses’, the event type (chasing or competition), and their interaction as fixed effects, with mouse identity as a random intercept. Model assumptions were tested using residual diagnostics and normality tests (Lilliefors, Anderson–Darling, and Shapiro–Wilk). Due to assumption violations, we performed cluster-resampled bootstrapping (10,000 iterations, resampled by subject) to estimate robust confidence intervals for fixed effects. The interaction term captured differences in the win–loss relationship between tube competitions and chasing events.

#### Relationship between social rank and chasing

First, to assess how tube competition metrics and chasing relate at the individual level, we conducted correlation analyses using Spearman’s rank correlation coefficients (cf. **Fig. 6A** and **Fig. 6C**). To account for repeated measures across mice, we also fitted LMEs with mouse identity as a random intercept that confirmed all results (see **Extended Data Main**). To further examine and illustrate structural patterns within social interactions, we processed win–loss match matrices from tube competition and chasing events for each NoSeMaze group. These match matrices were sorted by the David’s score (competition-based) or by the fraction of active chases (chasing-based) and normalized by dividing all entries by the maximum value within that matrix, thus setting the strongest observed interaction to 1. This procedure was applied at the group level to preserve the relative structure of interactions while allowing comparison across groups with differing activity levels. Normalized matrices were then averaged across all groups and visualized as heatmaps, with rows and columns representing individuals ordered by their social rank or active chases (cf. **Fig. 6D-F**). Color intensity indicated the relative frequency of interactions between each pair of individuals. To quantify asymmetries in interaction patterns, we modeled quadrant-level differences in the frequencies of normalized dyadic interactions using an LME with a beta distribution and logit link function. Proportions of dyadic events were bounded away from 0 and 1 by a small constant (ε = 0.0001) to meet the assumptions of the beta distribution. The model included quadrant as a fixed effect and group identity as a random intercept. Model fitting was performed using the glmmTMB package in R (https://cran.r-project.org/web/packages/glmmTMB/index.html) ^60^. Estimated marginal means were computed for each quadrant using the emmeans package (https://cran.r-project.org/web/packages/emmeans/index.html) ^61^, and back-transformed to the original scale (0–1) for interpretability. Pairwise post hoc comparisons between quadrants were conducted on the response scale, with Tukey-adjusted p-values to control for multiple comparisons.

#### Group-level analysis of social rank–chasing relationships

To evaluate the relationship between social rank and chasing at the group level, we computed Spearman’s correlation coefficients between the David’s scores from tube competitions and the fraction of active chases within each NoSeMaze group. This yielded one correlation coefficient per group (n = 18). We then tested whether these group-level coefficients were associated with group-level transitivity scores, using a Spearman’s rank correlation across groups (cf. **Fig. 6B**). This analysis assessed whether the strength of the within-group association between social rank and active chases varied systematically with the clarity of the social hierarchy. To further characterize this relationship, we also correlated group transitivity with the fraction of active chases performed by the mouse with the highest social rank (cf. **Supplementary Fig. S11**), reflecting the extent to which dominant individuals engaged in proactive aggression as a function of group hierarchy structure.

#### Relationships between social and reward-seeking features

Differences in the David’s score from tube competitions, the fraction of active chases and the fraction of times being chased between the three learning clusters were similarly assessed as for the six stimulus-outcome learning features (cf. **Clustering of reward-seeking profiles**), using the same median-based permutation method accounting for between- and within-animal variability. Again, to control for multiple comparisons, p-values were adjusted using the Benjamini–Hochberg FDR procedure ^54^. Then, association between the social metrics (David’s score and active chases) and the reward-seeking metrics (correct hit and rejection rate, pre-CS licking rate, CS+ modulation peak, switch latencies for CS+ and CS− from reversal 1 to 4 and for the median) were assessed with similar Spearman’s correlation analyses and LMEs, as described above (cf. **Relationship between social hierarchy and chasing**). Lastly, we performed principal component analysis (PCA) on z-scored data to examine the underlying structure of social and cognitive features more profoundly. Skewed variables were first log- or square root-transformed based on skewness direction and magnitude. PCA was conducted combining social (David’s score, active chases and being chased) and reward-seeking features. Loadings were interpreted to assess the contribution of each metric to the top three components. To evaluate domain specificity, permutation tests (n = 10,000) were used to test whether social and cognitive metrics differed significantly in their contributions to each principal component.

#### Genotype effects

Group differences in social and reinforcement-learning metrics (cf. **Supplementary Fig. S9** and **Supplementary Fig. S10**) between OXTR^ΔAON^ and control animals were assessed using permutation tests (10,000 iterations) comparing group medians. To account for repeated measures, group labels were shuffled at the level of the animal. Week-wise comparisons for the social metrics were FDR-corrected for multiple comparisons. To test whether the observed behavioral associations were influenced by the OXTR^ΔAON^ genotype, genotype (OXTR^ΔAON^ vs. control) was included as a covariate in LMEs and correlation analyses (see **Extended Data Main** and **Extended Data Supplement**). Specifically, LMEs were extended with the covariate as additional predictor, and Spearman’s partial correlation analyses were performed including the covariate as factor of no interest. In each analysis, we tested whether the primary relationships of interest remained significant when accounting for genotype. All analyses remained statistically significant after inclusion of the genotype covariate, indicating that the observed associations were robust across genotypes. For instance, in the relationship between the David’s score from tube competitions and the fraction of active chases, the inclusion of genotype in the LME did not alter the significance of the main association.

## Supporting information

Supplementary Material - Systematic Statistical Reporting

## Software and Data Availability

All behavioral data and analysis scripts used in this study are publicly available on GitHub: https://github.com/KelschLAB/NoSeMaze-social-status. The majority of the analyses were conducted in MATLAB, including the detection of behavioral events, the preprocessing, most of the analyses and the clustering procedures. Elo ratings and hierarchy structure assessments were performed using R (v4.4.1) and RStudio, employing the aniDom, EloRating, and lme4 packages. The NoSeMaze infrastructure was controlled by a modified version of the open-source Python-based software originally developed by Erskine et al. (2019), fully documented in the repository https://github.com/KelschLAB/NoSeMaze. All custom scripts and data necessary to reproduce the findings of this study are available upon request or via the linked repository.

## Data and Code Availability

The full documentation of the NoSeMaze, including hardware information and the open-source Python-based software package, is available on GitHub: https://github.com/KelschLAB/NoSeMaze. Source data files for each main figure and each supplementary figures as well as the extended data are provided with the manuscript. All analysis code and data matrices used to generate the results are available on GitHub: https://github.com/KelschLAB/NoSeMaze-social-status.

## Acknowledgements

We thank Michael Bram and Jan Ringkamp for their help in engineering the NoSeMaze, and Dr. Carla Filosa for her initial help in developing the behavioral analyses. We also thank Michal Adveev and Laura Haenschke for help with preparing the competition videos.

We acknowledge the use of ChatGPT (OpenAI, GPT-4, accessed 2025) for assistance with language editing, improving readability, and annotating code. All outputs were reviewed, validated, and verified by the authors, who take full responsibility for the content of this manuscript. The work was funded by BMBF 3R consortium grants ‘NoSeMaze1’ (161L0277A) and ‘NoSeMaze2’ (16LW0333K) to W.K., Leibniz Association program grant ‘Learning resilience’ (K430/2021) to W.K., Boehringer Ingelheim Foundation grant ‘Complex Systems’ to W.K., BMBF CRCNS grant ‘Oxystate’ (01GQ1708) to W.K, DFG CRC 379 Project C03 to W.K., and the DFG Clinician Scientist Program ‘Interfaces and Interventions in Complex Chronic Conditions’ (EB187/8-1) to J.R.

## References

1. Chase, I.D. (1980). Social process and hierarchy formation in small groups: a comparative perspective. American Sociological Review, 905–924.

2. Williamson, C.M., Lee, W., and Curley, J.P. (2016). Temporal dynamics of social hierarchy formation and maintenance in male mice. Animal Behaviour 115, 259–272. 10.1016/j.anbehav.2016.03.004.

3. Tibbetts, E.A., Pardo-Sanchez, J., and Weise, C. (2022). The establishment and maintenance of dominance hierarchies. Philosophical Transactions of the Royal Society B 377, 20200450.

4. Beery, A.K., and Kaufer, D. (2015). Stress, social behavior, and resilience: insights from rodents. Neurobiology of stress 1, 116–127.

5. LeClair, K.B., Chan, K.L., Kaster, M.P., Parise, L.F., Burnett, C.J., and Russo, S.J. (2021). Individual history of winning and hierarchy landscape influence stress susceptibility in mice. eLife 10, e71401. 10.7554/eLife.71401.

6. Sapolsky, R.M. (2005). The influence of social hierarchy on primate health. Science (New York, N.Y.) 308, 648–652.

7. Cum, M., Santiago Pérez, J.A., Iwata, R.L., Lopez, N., Higgs, A., Li, A., Ye, C., Wangia, E., Wright, E.S., García Restrepo, C., et al. (2024). A Multiparadigm Approach to Characterize Dominance Behaviors in CD1 and C57BL6 Male Mice. eneuro *11*, ENEURO.0342–0324.2024. 10.1523/eneuro.0342-24.2024.

8. Weber, E.M., Zidar, J., Ewaldsson, B., Askevik, K., Udén, E., Svensk, E., and Törnqvist, E. (2022). Aggression in Group-Housed Male Mice: A Systematic Review. Animals (Basel) 13. 10.3390/ani13010143.

9. Fulenwider, H.D., Caruso, M.A., and Ryabinin, A.E. (2022). Manifestations of domination: Assessments of social dominance in rodents. Genes Brain Behav 21, e12731. 10.1111/gbb.12731.

10. Lee, W., Khan, A., and Curley, J.P. (2017). Major urinary protein levels are associated with social status and context in mouse social hierarchies. Proceedings of the Royal Society B: Biological Sciences 284, 20171570.

11. Winiarski, M., Madecka, A., Yadav, A., Borowska, J., Wołyniak, M.R., Jędrzejewska-Szmek, J., Kondrakiewicz, L., Mankiewicz, L., Chaturvedi, M., Wójcik, D.K., et al. (2025). Information sharing within a social network is key to behavioral flexibility—Lessons from mice tested under seminaturalistic conditions. Science Advances 11, eadm7255. doi:10.1126/sciadv.adm7255.

12. Shemesh, Y., Sztainberg, Y., Forkosh, O., Shlapobersky, T., Chen, A., and Schneidman, E. (2013). High-order social interactions in groups of mice. eLife 2, e00759. 10.7554/eLife.00759.

13. Forkosh, O., Karamihalev, S., Roeh, S., Alon, U., Anpilov, S., Touma, C., Nussbaumer, M., Flachskamm, C., Kaplick, P.M., Shemesh, Y., et al. (2019). Identity domains capture individual differences from across the behavioral repertoire. Nature neuroscience 22, 2023–2028. 10.1038/s41593-019-0516-y.

14. Varholick, J.A., Bailoo, J.D., Jenkins, A., Voelkl, B., and Würbel, H. (2021). A Systematic Review and Meta-Analysis of the Relationship Between Social Dominance Status and Common Behavioral Phenotypes in Male Laboratory Mice. Frontiers in Behavioral Neuroscience 14. 10.3389/fnbeh.2020.624036.

15. Jupp, B., Murray, J.E., Jordan, E.R., Xia, J., Fluharty, M., Shrestha, S., Robbins, T.W., and Dalley, J.W. (2016). Social dominance in rats: effects on cocaine self-administration, novelty reactivity and dopamine receptor binding and content in the striatum. Psychopharmacology 233, 579–589.

16. Sánchez-Salvador, L., Prados-Pardo, Á., Martín-González, E., Olmedo-Córdoba, M., Mora, S., and Moreno, M. (2021). The role of social stress in the development of inhibitory control deficit: a systematic review in preclinical models. International Journal of Environmental Research and Public Health 18, 4953.

17. Kahnau, P., Mieske, P., Wilzopolski, J., Kalliokoski, O., Mandillo, S., Hölter, S.M., Voikar, V., Amfim, A., Badurek, S., Bartelik, A., et al. (2023). A systematic review of the development and application of home cage monitoring in laboratory mice and rats. BMC biology 21, 256.

18. Crabbe, J.C., Wahlsten, D., and Dudek, B.C. (1999). Genetics of mouse behavior: interactions with laboratory environment. Science (New York, N.Y.) 284, 1670–1672. 10.1126/science.284.5420.1670.

19. Voikar, V., and Gaburro, S. (2020). Three Pillars of Automated Home-Cage Phenotyping of Mice: Novel Findings, Refinement, and Reproducibility Based on Literature and Experience. Front Behav Neurosci 14, 575434. 10.3389/fnbeh.2020.575434.

20. Chesler, E.J., Wilson, S.G., Lariviere, W.R., Rodriguez-Zas, S.L., and Mogil, J.S. (2002). Influences of laboratory environment on behavior. Nature neuroscience 5, 1101–1102. 10.1038/nn1102-1101.

21. Fan, Z., Zhu, H., Zhou, T., Wang, S., Wu, Y., and Hu, H. (2019). Using the tube test to measure social hierarchy in mice. Nature Protocols 14, 819–831.

22. Mertens, S., Vogt, M.A., Gass, P., Palme, R., Hiebl, B., and Chourbaji, S. (2019). Effect of three different forms of handling on the variation of aggression-associated parameters in individually and group-housed male C57BL/6NCrl mice. PloS one 14, e0215367.

23. Gouveia, K., and Hurst, J.L. (2017). Optimising reliability of mouse performance in behavioural testing: the major role of non-aversive handling. Scientific reports 7, 44999. 10.1038/srep44999.

24. Weissbrod, A., Shapiro, A., Vasserman, G., Edry, L., Dayan, M., Yitzhaky, A., Hertzberg, L., Feinerman, O., and Kimchi, T. (2013). Automated long-term tracking and social behavioural phenotyping of animal colonies within a semi-natural environment. Nature communications 4, 2018.

25. Sofer, Y., Zilkha, N., Gimpel, E., Wagner, S., Chuartzman, S.G., and Kimchi, T. (2024). Sexually dimorphic oxytocin circuits drive intragroup social conflict and aggression in wild house mice. Nature neuroscience 27, 1565–1573.

26. Barabas, A.J., Lucas, J.R., Erasmus, M.A., Cheng, H.W., and Gaskill, B.N. (2021). Who’s the Boss? Assessing Convergent Validity of Aggression Based Dominance Measures in Male Laboratory Mice, Mus Musculus. Front Vet Sci 8, 695948. 10.3389/fvets.2021.695948.

27. Williamson, C.M., Lee, W., Romeo, R.D., and Curley, J.P. (2017). Social context-dependent relationships between mouse dominance rank and plasma hormone levels. Physiology & behavior 171, 110–119.

28. Chase, I.D., Tovey, C., Spangler-Martin, D., and Manfredonia, M. (2002). Individual differences versus social dynamics in the formation of animal dominance hierarchies. Proceedings of the National Academy of Sciences 99, 5744–5749.

29. Bengston, S.E., and Jandt, J.M. (2014). The development of collective personality: the ontogenetic drivers of behavioral variation across groups. Frontiers in Ecology and Evolution 2, 81.

30. Freund, J., Brandmaier, A.M., Lewejohann, L., Kirste, I., Kritzler, M., Krüger, A., Sachser, N., Lindenberger, U., and Kempermann, G. (2013). Emergence of individuality in genetically identical mice. Science (New York, N.Y.) 340, 756–759.

31. Cantor, M., Maldonado-Chaparro, A.A., Beck, K.B., Brandl, H.B., Carter, G.G., He, P., Hillemann, F., Klarevas-Irby, J.A., Ogino, M., Papageorgiou, D., et al. (2021). The importance of individual-to-society feedbacks in animal ecology and evolution. Journal of Animal Ecology 90, 27–44.

32. Oettl, L.-L., Ravi, N., Schneider, M., Scheller, Max F., Schneider, P., Mitre, M., da Silva Gouveia, M., Froemke, Robert C., Chao, Moses V., Young, W.S., et al. (2016). Oxytocin Enhances Social Recognition by Modulating Cortical Control of Early Olfactory Processing. Neuron 90, 609–621. 10.1016/j.neuron.2016.03.033.

33. Wolf, D., Hartig, R., Zhuo, Y., Scheller, M.F., Articus, M., Moor, M., Grinevich, V., Linster, C., Russo, E., Weber-Fahr, W., et al. (2024). Oxytocin induces the formation of distinctive cortical representations and cognitions biased toward familiar mice. Nature Communications 15, 6274. 10.1038/s41467-024-50113-6.

34. Bari, A., and Robbins, T.W. (2013). Inhibition and impulsivity: behavioral and neural basis of response control. Progress in neurobiology 108, 44–79.

35. Sánchez-Tójar, A., Schroeder, J., and Farine, D.R. (2018). A practical guide for inferring reliable dominance hierarchies and estimating their uncertainty. Journal of Animal Ecology 87, 594–608.

36. Nelias, C., Ghanayem, S., Wolf, D., Moor, M., Scheller, M.F., Grinevich, V., Reinwald, J.R., and Kelsch, W. (2025). Stable clique membership in mouse societies requires oxytocin-enabled social sensory states. bioRxiv. 10.1101/2025.08.26.672298.

37. Bernstein, I.S. (1980). Dominance: A theoretical perspective for ethologists. In Dominance relations: An ethological view of human conflict and social interaction, D.S. Omark, FF; Freedman, DG;, ed. (Garland Press: New York), pp. 71–84.

38. Torquet, N., Marti, F., Campart, C., Tolu, S., Nguyen, C., Oberto, V., Benallaoua, M., Naudé, J., Didienne, S., Debray, N., et al. (2018). Social interactions impact on the dopaminergic system and drive individuality. Nature communications 9, 3081.

39. Castelhano-Carlos, M., Costa, P.S., Russig, H., and Sousa, N. (2014). PhenoWorld: a new paradigm to screen rodent behavior. Translational psychiatry 4, e399–e399.

40. Erskine, A., Bus, T., Herb, J.T., and Schaefer, A.T. (2019). AutonoMouse: High throughput operant conditioning reveals progressive impairment with graded olfactory bulb lesions. PloS one 14, e0211571.

41. Kiryk, A., Janusz, A., Zglinicki, B., Turkes, E., Knapska, E., Konopka, W., Lipp, H.-P., and Kaczmarek, L. (2020). IntelliCage as a tool for measuring mouse behavior–20 years perspective. Behavioural brain research 388, 112620.

42. Lindzey, G., Winston, H., and Manosevitz, M. (1961). Social dominance in inbred mouse strains.

43. Wang, F., Zhu, J., Zhu, H., Zhang, Q., Lin, Z., and Hu, H. (2011). Bidirectional control of social hierarchy by synaptic efficacy in medial prefrontal cortex. Science (New York, N.Y.) 334, 693–697. 10.1126/science.1209951.

44. Miczek, K.A., Maxson, S.C., Fish, E.W., and Faccidomo, S. (2001). Aggressive behavioral phenotypes in mice. Behavioural brain research 125, 167–181.

45. Clutton-Brock, T.H., and Huchard, E. (2013). Social competition and selection in males and females. Philosophical Transactions of the Royal Society B: Biological Sciences 368, 20130074.

46. Van Loo, P.L., Mol, J.A., Koolhaas, J.M., Van Zutphen, B.F., and Baumans, V. (2001). Modulation of aggression in male mice: influence of group size and cage size. Physiology & behavior 72, 675–683.

47. Cervantes, M.C., Laughlin, R.E., and Jentsch, J.D. (2013). Cocaine self-administration behavior in inbred mouse lines segregating different capacities for inhibitory control. Psychopharmacology 229, 515–525.

48. Dongelmans, M., Durand-de Cuttoli, R., Nguyen, C., Come, M., Duranté, E.K., Lemoine, D., Brito, R., Ahmed Yahia, T., Mondoloni, S., Didienne, S., et al. (2021). Chronic nicotine increases midbrain dopamine neuron activity and biases individual strategies towards reduced exploration in mice. Nature Communications 12, 6945.

49. Huang, Y., Luan, S., Wu, B., Li, Y., Wu, J., Chen, W., and Hertwig, R. (2024). Impulsivity is a stable, measurable, and predictive psychological trait. Proceedings of the National Academy of Sciences 121, e2321758121.

50. Pittaras, E., Callebert, J., Chennaoui, M., Rabat, A., and Granon, S. (2016). Individual behavioral and neurochemical markers of unadapted decision-making processes in healthy inbred mice. Brain Structure and Function 221, 4615–4629.

51. Magnus, C.J., Lee, P.H., Bonaventura, J., Zemla, R., Gomez, J.L., Ramirez, M.H., Hu, X., Galvan, A., Basu, J., Michaelides, M., et al. (2019). Ultrapotent chemogenetics for research and potential clinical applications. Science (New York, N.Y.) 364, eaav5282.

52. Tost, H., Reichert, M., Braun, U., Reinhard, I., Peters, R., Lautenbach, S., Hoell, A., Schwarz, E., Ebner-Priemer, U., Zipf, A., et al. (2019). Neural correlates of individual differences in affective benefit of real-life urban green space exposure. Nature neuroscience 22, 1389–1393. 10.1038/s41593-019-0451-y.

53. Reichert, M., Gan, G., Renz, M., Braun, U., Brüßler, S., Timm, I., Ma, R., Berhe, O., Benedyk, A., Moldavski, A., et al. (2021). Ambulatory assessment for precision psychiatry: Foundations, current developments and future avenues. Experimental Neurology 345, 113807. 10.1016/j.expneurol.2021.113807.

54. Benjamini, Y., and Hochberg, Y. (1995). Controlling the false discovery rate: a practical and powerful approach to multiple testing. Journal of the Royal statistical society: series B (Methodological) 57, 289–300.

55. David, H.A. (1987). Ranking from unbalanced paired-comparison data. Biometrika 74, 432–436.

56. Shizuka, D., and McDonald, D.B. (2012). A social network perspective on measurements of dominance hierarchies. Animal Behaviour 83, 925–934. 10.1016/j.anbehav.2012.01.011.

57. de Vries, H., Stevens, J.M.G., and Vervaecke, H. (2006). Measuring and testing the steepness of dominance hierarchies. Animal Behaviour 71, 585–592. 10.1016/j.anbehav.2005.05.015.

58. Neumann, C., Duboscq, J., Dubuc, C., Ginting, A., Irwan, A.M., Agil, M., Widdig, A., and Engelhardt, A. (2011). Assessing dominance hierarchies: validation and advantages of progressive evaluation with Elo-rating. Animal Behaviour 82, 911–921.

59. Bates, D., Mächler, M., Bolker, B., and Walker, S. (2015). Fitting linear mixed-effects models using lme4. Journal of statistical software 67, 1–48.

60. Brooks, M.E., Kristensen, K., Van Benthem, K.J., Magnusson, A., Berg, C.W., Nielsen, A., Skaug, H.J., Mächler, M., and Bolker, B.M. (2017). glmmTMB balances speed and flexibility among packages for zero-inflated generalized linear mixed modeling.

61. Lenth, R. (2023). emmeans: Estimated Marginal Means, aka Least-Squares Means_. R package version 1.8. 5.

