## Supplementary Material for "Individual differences drive social hierarchies in male mouse societies"

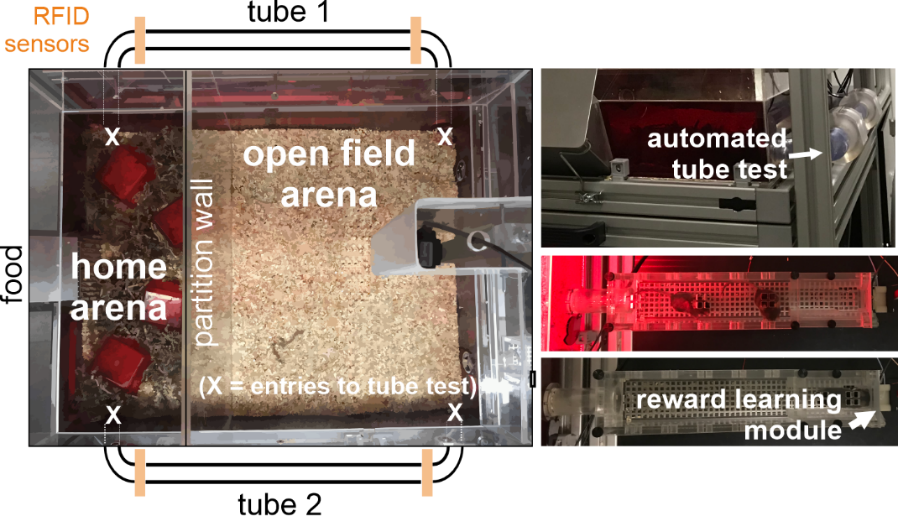

**Fig. S1: Illustration of NoSeMaze modules.**

Illustration of the NoSeMaze containing the automated tube test and the reward learning module.

*NoSeMaze, non-invasive sensor rich maze; RFID, radio-frequency identification*

| **Mouse RFID** | **group** | **repetition** | **age** | **genotype** | **weight** | **Mouse RFID** | **group** | **repetition** | **age** | **genotype** | **weight** |
| --- | --- | --- | --- | --- | --- | --- | --- | --- | --- | --- | --- |
| 0007AC2B02 | 1 | 1 | 55 | WT | 33.59 | 0007CB0A48 | 12 | 1 | 18 | WT | 28.38 |
| 0007CB0942 | 1 | 1 | 55 | WT | 34.36 | 0007CB0AD4 | 12 | 1 | 16 | WT | 23.28 |
| 0007CB44CE | 1 | 1 | 55 | WT | 25.92 | 0007CB2ED2 | 12 | 1 | 16 | WT | 26.51 |
| 0007CB7051 | 1 | 1 | 56 | WT | 32.58 | 0007CB321B | 12 | 1 | 18 | WT | 26.61 |
| 0007CD6778 | 1 | 1 | 57 | OXTR^ΔAON^ | 36.42 | 0007CB39C9 | 12 | 1 | 18 | OXTR^ΔAON^ | 23.49 |
| 0007CD765E | 1 | 1 | 57 | WT | 31.68 | 0007CB464C | 12 | 1 | 16 | OXTR^ΔAON^ | 27.25 |
| 0007CD77C5 | 1 | 1 | 57 | OXTR^ΔAON^ | 32.71 | 0007F2B770 | 12 | 1 | 17 | OXTR^ΔAON^ | 27.57 |
| 0007CDADED | 1 | 1 | 55 | WT | 36.89 | 0007F2B957 | 12 | 1 | 18 | OXTR^ΔAON^ | 28.45 |
| 0007CDC443 | 1 | 1 | 55 | WT | 36.46 | 0007F2BA9B | 12 | 1 | 18 | WT | 26.04 |
| 0007CDDBAF | 1 | 1 | 57 | WT | 31.46 | 0007F2CA05 | 12 | 1 | 18 | OXTR^ΔAON^ | 25.31 |
| 0007CB2224 | 2 | 1 | 58 | WT | 35.99 | 0007CA3B1C | 13 | 1 | 17 | WT | 24.43 |
| 0007CB3491 | 2 | 1 | 56 | WT | 33.91 | 0007CB1E1F | 13 | 1 | 18 | WT | 28.15 |
| 0007CB4837 | 2 | 1 | 56 | OXTR^ΔAON^ | 40.03 | 0007CB2183 | 13 | 1 | 17 | OXTR^ΔAON^ | 25.32 |
| 0007CB6AB4 | 2 | 1 | 56 | WT | 27.19 | 0007CB2277 | 13 | 1 | 18 | OXTR^ΔAON^ | 26.24 |
| 0007CD5116 | 2 | 1 | 57 | WT | 36.69 | 0007CB3D9B | 13 | 1 | 18 | WT | 24.12 |
| 0007CDE223 | 2 | 1 | 75 | WT | 33.16 | 0007CB3F5E | 13 | 1 | 18 | WT | 25.26 |
| 0007CDE4F0 | 2 | 1 | 58 | OXTR^ΔAON^ | 30.71 | 0007CB6BC3 | 13 | 1 | 16 | WT | 23.81 |
| 0007CDEAA7 | 2 | 1 | 57 | WT | 36.04 | 0007F2945F | 13 | 1 | 18 | WT | 23.81 |
| 0007CDEC0B | 2 | 1 | 75 | WT | 32.32 | 0007F2B8A6 | 13 | 1 | 17 | OXTR^ΔAON^ | 24.56 |
| 0007CDEEE2 | 2 | 1 | 75 | WT | 32.92 | 0007F2E455 | 13 | 1 | 18 | OXTR^ΔAON^ | 24.72 |
| 0007CB0942 | 3 | 2 | 59 | WT | 34.47 | 0007CB0CDD | 14 | 1 | 16 | WT | 23.82 |
| 0007CD38E5 | 3 | 1 | 59 | WT | 35.51 | 0007CB0F67 | 14 | 1 | 16 | OXTR^ΔAON^ | 27.03 |
| 0007CD7FB2 | 3 | 1 | 63 | OXTR^ΔAON^ | 34.07 | 0007CB48B4 | 14 | 1 | 16 | WT | 27.09 |
| 0007CD81D3 | 3 | 1 | 59 | WT | 37.51 | 0007CB6DFA | 14 | 1 | 16 | OXTR^ΔAON^ | 24.81 |
| 0007CD86B3 | 3 | 1 | 78 | WT | 32.71 | 0007F29741 | 14 | 1 | 17 | WT | 25.78 |
| 0007CDAD6B | 3 | 1 | 78 | WT | 28.63 | 0007F2B88A | 14 | 1 | 17 | WT | 24.38 |
| 0007CDC395 | 3 | 1 | 63 | OXTR^ΔAON^ | 34.19 | 0007F2C687 | 14 | 1 | 18 | WT | 26.08 |
| 0007CDC91B | 3 | 1 | 78 | WT | 31.69 | 0007F2EEFA | 14 | 1 | 17 | WT | 24.78 |
| 0007CDDBAF | 3 | 2 | 61 | WT | 32.41 | 0007F2F26A | 14 | 1 | 18 | WT | 25.74 |
| 0007CE0327 | 3 | 1 | 78 | WT | 29.83 | 0007F2F283 | 14 | 1 | 16 | OXTR^ΔAON^ | 24.07 |
| 0007ABF555 | 4 | 1 | 62 | OXTR^ΔAON^ | 32.48 | 0007CA3A72 | 15 | 2 | 22 | OXTR^ΔAON^ | 26.26 |
| 0007AC2B02 | 4 | 2 | 60 | WT | 33.03 | 0007CB0B74 | 15 | 2 | 22 | WT | 27.02 |
| 0007ACA781 | 4 | 1 | 79 | WT | 32.69 | 0007CB0DBC | 15 | 2 | 22 | OXTR^ΔAON^ | 28.11 |
| 0007CB2224 | 4 | 2 | 62 | WT | 35.46 | 0007CB2407 | 15 | 2 | 22 | OXTR^ΔAON^ | 24.43 |
| 0007CB7051 | 4 | 2 | 61 | WT | 32.78 | 0007F2B3FF | 15 | 2 | 22 | WT | 26.18 |
| 0007CD6778 | 4 | 2 | 62 | OXTR^ΔAON^ | 38.56 | 0007F2B59E | 15 | 2 | 20 | WT | 28.52 |
| 0007CD765E | 4 | 2 | 62 | WT | 30.89 | 0007F2B8A2 | 15 | 2 | 22 | OXTR^ΔAON^ | 26.42 |
| 0007CDADED | 4 | 2 | 60 | WT | 37.41 | 0007F2B9FC | 15 | 2 | 22 | WT | 26.52 |
| 0007CDC443 | 4 | 2 | 60 | WT | 37.22 | 0007F2E37E | 15 | 2 | 22 | WT | 28.04 |
| 0007CDEAA7 | 4 | 2 | 60 | WT | 35.07 | 0007F2E3D4 | 15 | 2 | 22 | WT | 26.56 |
| 0007CB0942 | 5 | 3 | 62 | WT | 34.51 | 0007CA3B1C | 16 | 2 | 21 | WT | 26.49 |
| 0007CB3491 | 5 | 2 | 62 | WT | 32.79 | 0007CB1E1F | 16 | 2 | 22 | WT | 28.75 |
| 0007CB4837 | 5 | 2 | 62 | OXTR^ΔAON^ | 36.62 | 0007CB2183 | 16 | 2 | 21 | OXTR^ΔAON^ | 25.83 |
| 0007CD72E3 | 5 | 1 | 81 | WT | 35.94 | 0007CB2277 | 16 | 2 | 22 | OXTR^ΔAON^ | 27.41 |
| 0007CD77C5 | 5 | 2 | 64 | OXTR^ΔAON^ | 32.59 | 0007CB3D9B | 16 | 2 | 22 | WT | 25.72 |
| 0007CDDBAF | 5 | 3 | 64 | WT | 31.31 | 0007CB3F5E | 16 | 2 | 22 | WT | 26.67 |
| 0007CDE223 | 5 | 2 | 81 | WT | 31.39 | 0007CB6BC3 | 16 | 2 | 20 | WT | 26.87 |
| 0007CDEC0B | 5 | 2 | 81 | WT | 31.29 | 0007F2945F | 16 | 2 | 22 | WT | 24.34 |
| 0007CDEEE2 | 5 | 2 | 81 | WT | 31.82 | 0007F2B8A6 | 16 | 2 | 21 | OXTR^ΔAON^ | 26.76 |
| 0007CE0327 | 5 | 2 | 81 | WT | 28.41 | 0007F2E455 | 16 | 2 | 22 | OXTR^ΔAON^ | 25.24 |
| 0007CB6AB4 | 6 | 2 | 73 | WT | 29.18 | 0007CB0CDD | 17 | 2 | 20 | WT | 25.19 |
| 0007CB7051 | 6 | 3 | 74 | WT | 33.63 | 0007CB0F67 | 17 | 2 | 20 | OXTR^ΔAON^ | 29.21 |
| 0007CD5116 | 6 | 3 | 74 | WT | 31.61 | 0007CB48B4 | 17 | 2 | 20 | WT | 26.15 |
| 0007CD765E | 6 | 3 | 75 | WT | 31.39 | 0007CB6DFA | 17 | 2 | 20 | OXTR^ΔAON^ | 25.86 |
| 0007CD86B3 | 6 | 3 | 91 | WT | 32.58 | 0007F29741 | 17 | 2 | 21 | WT | 25.49 |
| 0007CDAD6B | 6 | 2 | 91 | WT | 27.82 | 0007F2B88A | 17 | 2 | 21 | WT | 25.22 |
| 0007CDADED | 6 | 3 | 73 | WT | 36.98 | 0007F2C687 | 17 | 2 | 22 | WT | 25.96 |
| 0007CDC395 | 6 | 2 | 77 | OXTR^ΔAON^ | 32.67 | 0007F2EEFA | 17 | 2 | 21 | WT | 25.61 |
| 0007CDC443 | 6 | 3 | 73 | WT | 34.07 | 0007F2F26A | 17 | 2 | 22 | WT | 26.72 |
| 0007CDE4F0 | 6 | 2 | 75 | OXTR^ΔAON^ | 30.01 | 0007F2F283 | 17 | 2 | 20 | OXTR^ΔAON^ | 25.74 |
| 0007ABF555 | 7 | 2 | 68 | OXTR^ΔAON^ | 31.61 | 0007CA3B1C | 18 | 3 | 26 | WT | 27.78 |
| 0007ACA781 | 7 | 2 | 85 | WT | 31.38 | 0007CB0AD4 | 18 | 3 | 25 | WT | 23.92 |
| 0007CB2224 | 7 | 3 | 68 | WT | 35.72 | 0007CB0F67 | 18 | 3 | 25 | OXTR^ΔAON^ | 29.38 |
| 0007CD38E5 | 7 | 2 | 66 | WT | 34.85 | 0007CB2183 | 18 | 3 | 26 | OXTR^ΔAON^ | 26.82 |
| 0007CD72E3 | 7 | 2 | 85 | WT | 34.39 | 0007CB2407 | 18 | 3 | 27 | OXTR^ΔAON^ | 25.84 |
| 0007CD7FB2 | 7 | 2 | 70 | OXTR^ΔAON^ | 33.79 | 0007CB3F5E | 18 | 3 | 27 | WT | 29.41 |
| 0007CD81D3 | 7 | 2 | 66 | WT | 34.29 | 0007CB48B4 | 18 | 3 | 25 | WT | 27.93 |
| 0007CDEAA7 | 7 | 3 | 66 | WT | 34.37 | 0007F2BA9B | 18 | 3 | 27 | WT | 28.36 |
| 0007CDEEE2 | 7 | 3 | 85 | WT | 30.45 | 0007F2E3D4 | 18 | 3 | 27 | WT | 28.97 |
| 0007CE0327 | 7 | 3 | 85 | WT | 29.61 | 0007F2E455 | 18 | 3 | 27 | OXTR^ΔAON^ | 26.29 |
| 0007CA3A3A | 8 | 1 | 68 | WT | 25.47 | 0007CB0AD4 | 19 | 2 | 20 | WT | 22.81 |
| 0007CB0942 | 8 | 4 | 68 | WT | 34.24 | 0007CB2ED2 | 19 | 2 | 20 | WT | 30.42 |
| 0007CB30A6 | 8 | 1 | 72 | OXTR^ΔAON^ | 30.41 | 0007CB321B | 19 | 2 | 22 | WT | 26.19 |
| 0007CB486D | 8 | 1 | 71 | OXTR^ΔAON^ | 31.92 | 0007CB39C9 | 19 | 2 | 22 | OXTR^ΔAON^ | 26.18 |
| 0007CD5116 | 8 | 2 | 69 | WT | 32.67 | 0007F2B770 | 19 | 2 | 21 | OXTR^ΔAON^ | 27.57 |
| 0007CD86B3 | 8 | 2 | 87 | WT | 32.68 | 0007F2B957 | 19 | 2 | 22 | OXTR^ΔAON^ | 30.19 |
| 0007CDC91B | 8 | 2 | 87 | WT | 32.73 | 0007F2BA9B | 19 | 2 | 22 | WT | 26.18 |
| 0007CDDBAF | 8 | 4 | 70 | WT | 32.51 | 0007F2CA05 | 19 | 2 | 22 | OXTR^ΔAON^ | 25.88 |
| 0007CDE223 | 8 | 3 | 87 | WT | 33.11 | 0007F2E42C | 19 | 1 | 22 | WT | 30.91 |
| 0007CDEC0B | 8 | 3 | 87 | WT | 31.56 | 0007CB0DBC | 20 | 3 | 27 | OXTR^ΔAON^ | 30.82 |
| 0007CB2224 | 9 | 4 | 73 | WT | 36.15 | 0007CB1E1F | 20 | 3 | 27 | WT | 30.65 |
| 0007CB2302 | 9 | 1 | 71 | OXTR^ΔAON^ | 28.67 | 0007CB321B | 20 | 3 | 27 | WT | 28.21 |
| 0007CB3713 | 9 | 1 | 71 | WT | 32.15 | 0007F29741 | 20 | 3 | 26 | WT | 27.97 |
| 0007CB40DE | 9 | 1 | 71 | WT | 25.89 | 0007F2B770 | 20 | 3 | 26 | OXTR^ΔAON^ | 29.14 |
| 0007CB47A6 | 9 | 1 | 72 | WT | 36.48 | 0007F2B9FC | 20 | 3 | 27 | WT | 27.96 |
| 0007CB486D | 9 | 2 | 74 | OXTR^ΔAON^ | 30.16 | 0007F2CA05 | 20 | 3 | 27 | OXTR^ΔAON^ | 26.27 |
| 0007CDEC0B | 9 | 4 | 90 | WT | 31.29 | 0007F2E37E | 20 | 3 | 27 | WT | 29.92 |
| 0007CDEEE2 | 9 | 4 | 90 | WT | 31.51 | 0007F2EEFA | 20 | 3 | 26 | WT | 28.03 |
| 0007CE0327 | 9 | 4 | 90 | WT | 29.14 | 0007F2F283 | 20 | 3 | 25 | OXTR^ΔAON^ | 27.64 |
| 0007CA3A3A | 10 | 2 | 75 | WT | 25.77 | 0007CA3A72 | 21 | 3 | 27 | OXTR^ΔAON^ | 27.51 |
| 0007CB0942 | 10 | 5 | 75 | WT | 35.77 | 0007CB2277 | 21 | 3 | 27 | OXTR^ΔAON^ | 29.32 |
| 0007CB2302 | 10 | 2 | 75 | OXTR^ΔAON^ | 28.45 | 0007CB464C | 21 | 3 | 25 | OXTR^ΔAON^ | 28.02 |
| 0007CB30A6 | 10 | 2 | 79 | OXTR^ΔAON^ | 29.28 | 0007F2B3FF | 21 | 3 | 27 | WT | 27.05 |
| 0007CB3713 | 10 | 2 | 75 | WT | 31.29 | 0007F2B59E | 21 | 3 | 25 | WT | 31.52 |
| 0007CB47A6 | 10 | 2 | 76 | WT | 35.97 | 0007F2B88A | 21 | 3 | 26 | WT | 28.08 |
| 0007CDDBAF | 10 | 5 | 77 | WT | 31.39 | 0007F2B8A6 | 21 | 3 | 26 | OXTR^ΔAON^ | 28.69 |
| 0007CDE223 | 10 | 4 | 94 | WT | 32.49 | 0007F2B957 | 21 | 3 | 27 | OXTR^ΔAON^ | 31.32 |
| 0007CDEC0B | 10 | 5 | 94 | WT | 30.08 | 0007F2C687 | 21 | 3 | 27 | WT | 27.81 |
| 0007CE0327 | 10 | 5 | 94 | WT | 28.62 | 0007F2F26A | 21 | 3 | 27 | WT | 28.52 |
| 0007CA3A72 | 11 | 1 | 18 | OXTR^ΔAON^ | 26.68 |  |  |  |  |  |  |
| 0007CB0B74 | 11 | 1 | 18 | WT | 27.92 |  |  |  |  |  |  |
| 0007CB0DBC | 11 | 1 | 18 | OXTR^ΔAON^ | 28.07 |  |  |  |  |  |  |
| 0007CB2407 | 11 | 1 | 18 | OXTR^ΔAON^ | 24.12 |  |  |  |  |  |  |
| 0007F2B3FF | 11 | 1 | 18 | WT | 25.38 |  |  |  |  |  |  |
| 0007F2B59E | 11 | 1 | 16 | WT | 28.07 |  |  |  |  |  |  |
| 0007F2B8A2 | 11 | 1 | 18 | OXTR^ΔAON^ | 25.69 |  |  |  |  |  |  |
| 0007F2B9FC | 11 | 1 | 18 | WT | 25.71 |  |  |  |  |  |  |
| 0007F2E37E | 11 | 1 | 18 | WT | 26.1 |  |  |  |  |  |  |
| 0007F2E3D4 | 11 | 1 | 18 | WT | 26.16 |  |  |  |  |  |  |

**Table S1: Overview of group compositions and animal information.**

*OXTR^ΔAON^, bilateral oxytocin receptor deletion in the AON pars centralis; RFID, radio-frequency identification; WT, wild type*

| **group ID** | **n of mice** | **age**  **(median, SD, in weeks)** | **tube data** | **stimulus-outcome learning data** |
| --- | --- | --- | --- | --- |
| 1 | 10 | 55.5, 0.99 | acquired | acquired |
| 2 | 10 | 57.5, 8.79 | acquired | acquired |
| 3 | 10 | 63, 9.07 | acquired | acquired |
| 4 | 10 | 61.5, 5.77 | acquired | acquired |
| 5 | 10 | 72.5, 9.62 | acquired | acquired |
| 6 | 10 | 74.5, 7.17 | acquired | acquired |
| 7 | 10 | 69, 9.20 | acquired | acquired |
| 8 | 10 | 71.5, 9.03 | acquired | acquired |
| 9 | 9 | 73, 9.06 | acquired | acquired |
| 10 | 10 | 76.5, 8.78 | acquired | acquired |
| 11 | 10 | 18, 0.63 | acquired | acquired |
| 12 | 10 | 18, 0.95 | acquired | Reinforcement learning port not working |
| 13 | 10 | 18, 0.81 | acquired | Reinforcement learning port not working |
| 14 | 10 | 16.5, 0.82 | acquired | Reinforcement learning port not working |
| 15 | 10 | 22, 0.63 | acquired | Reinforcement learning port not working |
| 16 | 10 | 22, 0.71 | RFID detectors not working | acquired |
| 17 | 10 | 20.5, 0.82 | acquired | acquired |
| 18 | 10 | 26.5, 0.92 | acquired | acquired |
| 19 | 10 | 22, 0.88 | acquired | acquired |
| 20 | 10 | 27, 0.84 | RFID detectors not working | acquired |
| 21 | 10 | 27, 0.84 | RFID detectors not working | acquired |

**Table S2: Overview of group characteristics and acquisition of behavioral data in the NoSeMaze.**

*ID, identity; NoSeMaze, non-invasive sensor rich maze; STD, standard deviation*

| **Dataset** | **# rounds observed** | **n mice** | **%** |
| --- | --- | --- | --- |
| Tube (social rank/chasing) | 1 | 11 | 13.9% |
| Tube (social rank /chasing) | 2 | 48 | 60.8% |
| Tube (social rank /chasing) | 3 | 13 | 16.5% |
| Tube (social rank /chasing) | 4 | 3 | 3.8% |
| Tube (social rank /chasing) | 5 | 4 | 5.1% |
| Lickport (reinforcement learning) | 1 | 13 | 16.7% |
| Lickport (reinforcement learning) | 2 | 51 | 65.4% |
| Lickport (reinforcement learning) | 3 | 7 | 9.0% |
| Lickport (reinforcement learning) | 4 | 3 | 3.8% |
| Lickport (reinforcement learning) | 5 | 4 | 5.1% |

**Table S3: Round participation across datasets**

Distribution of the number of NoSeMaze rounds contributed per mouse for the tube dataset (competition-based social rank and chasing) and the lickport dataset (reinforcement-learning). Percentages refer to the total number of mice available in each dataset. Most animals contributed at least two rounds (tube: 68/79, 86.1%; lickport: 65/78, 83.3%), indicating that R1–R2 stability estimates are based on paired observations for the majority of subjects.

| **metric** | **n_pairs_ (R1–R2)** | **Spearman ρ (R1–R2)**  **[95% CI]** | **ICC**  **no age**  **[95% CI]** | **ICC**  **age-adj** | **ICC**  **balanced first2** |
| --- | --- | --- | --- | --- | --- |
| **Competition David’s score (z-scored)** | 68 | 0.57  [0.41, 0.69] | 0.55  [0.42, 0.69] | 0.55  [0.40, 0.68] | 0.56 |
| **Active chases**  **(fraction, cube root)** | 68 | 0.75  [0.55, 0.89] | 0.74  [0.58, 0.85] | 0.73  [0.58,0.84] | 0.72 |
| **Being chased**  **(fraction, cube root)** | 68 | 0.57  [0.35, 0.75] | 0.61  [0.47, 0.71] | 0.60  [0.45, 0.71] | 0.60 |
| **Hit rate**  **(boxcox)** | 62 | 0.34  [0.16, 0.53] | 0.21  [0.06, 0.36] | 0.23  [0.07, 0.40] | 0.21 |
| **Baseline lick rate**  **(boxcox)** | 62 | 0.69  [0.53, 0.81] | 0.40  [0.17, 0.66] | 0.35  [0.15, 0.61] | 0.59 |
| **Correct rejection rate**  **(boxcox)** | 62 | 0.78  [0.61, 0.88] | 0.65  [0.37, 0.79] | 0.68  [0.44, 0.82] | 0.80 |
| **CS+ modulation peak**  **(boxcoxs)** | 62 | 0.56  [0.37, 0.71] | 0.58  [0.34, 0.71] | 0.59  [0.34, 0.71] | 0.59 |
| **CS+ switch latency**  **(boxcox)** | 62 | 0.24  [0.00, 0.48] | 0.27  [0.13, 0.38] | 0.33  [0.16, 0.43] | 0.27 |
| **CS− switch latency**  **(boxcox)** | 61 | 0.03  [−0.26, 0.36] | 0.13  [<0.01, 0.09] | 0.10  [<0.01, 0.32] | 0.15 |

**Table S4. Cross-round stability of social and reinforcement-learning metrics.**

Stability was quantified by Spearman correlations between round 1 and round 2 (R1–R2) with 95% bootstrap confidence intervals, and intraclass correlation coefficients (ICCs) estimated from variance-component mixed-effects models across all available rounds. “ICC no age” denotes ICC_across_group_ from a REML model with mouse identity as a random intercept and group as a random effect (repetition included as a fixed effect, cf. Methods).

“ICC age-adj” includes an age decomposition (group-mean age and within-group deviation) as additional fixed effects. “ICC balanced first2” recomputes ICC_across_group_ in a conservative balanced subset restricted to each mouse’s first two observed sessions and mice with both sessions, to control for unequal round participation. n_pairs_ indicates the number of mice contributing data to both R1 and R2.

| **metric** | **repetition**  **(p_rep., cond._)** | **age block**  **(p_age, cond._)** | **β_mean_**  **(mean age per group)** | **p_mean_**  **(mean age per group)** | **β_rel_**  **(rel. age per group)** | **p_rel_**  **(rel. age per group)** |
| --- | --- | --- | --- | --- | --- | --- |
| **Competition David’s score** | **0.0472** | 0.6322 | -0.0024 | 0.5330 | 0.0109 | 0.5172 |
| **Active chases**  **(fraction, cube root)** | 0.4387 | **0.0040** | -0.0006 | 0.4139 | **0.0088** | **0.0021** |
| **Being chased**  **(fraction, cube root)** | 0.6056 | 0.0535 | -0.0001 | 0.7225 | **0.0040** | **0.0198** |
| **Hit rate**  **(boxcox)** | **0.0012** | **0.0239** | **-0.0004** | **0.0159** | 0.0006 | 0.1653 |
| **Baseline lick rate**  **(boxcox)** | 0.3514 | **0.0058** | **-0.0182** | **0.0102** | **-0.0408** | **0.0228** |
| **Correct rejection rate**  **(boxcox)** | 0.2973 | **0.0002** | **0.0039** | **<0.0001** | 0.0030 | 0.3124 |
| **CS+ modulation peak**  **(boxcoxs)** | 0.7783 | 0.9701 | 0.0123 | 0.8051 | 0.0054 | 0.9774 |
| **CS+ switch latency**  **(boxcox)** | 0.0843 | **0.0108** | **0.0283** | **0.0011** | -0.0011 | 0.9504 |
| **CS− switch latency**  **(boxcox)** | **0.0111** | 0.1581 | -0.0036 | 0.3200 | -0.0149 | 0.0810 |

**Table S5. Effects of repetition and age on mean metric levels.**

Mixed-effects model diagnostics testing whether metric values shift systematically with session experience (repetition) or age. Reported are likelihood-ratio test p-values for adding repetition conditional on the age block (p_rep, cond._) and for adding the age block conditional on repetition (p_age, cond._). The age block was decomposed into a between-group component (mean age per group; β_mean_) and a within-group component (each animal’s deviation from its group mean; β_rel_), with corresponding p-values for each coefficient from the full model. These tests assess mean-level shifts (age/experience effects) and are reported separately from the stability estimates in Table S4.

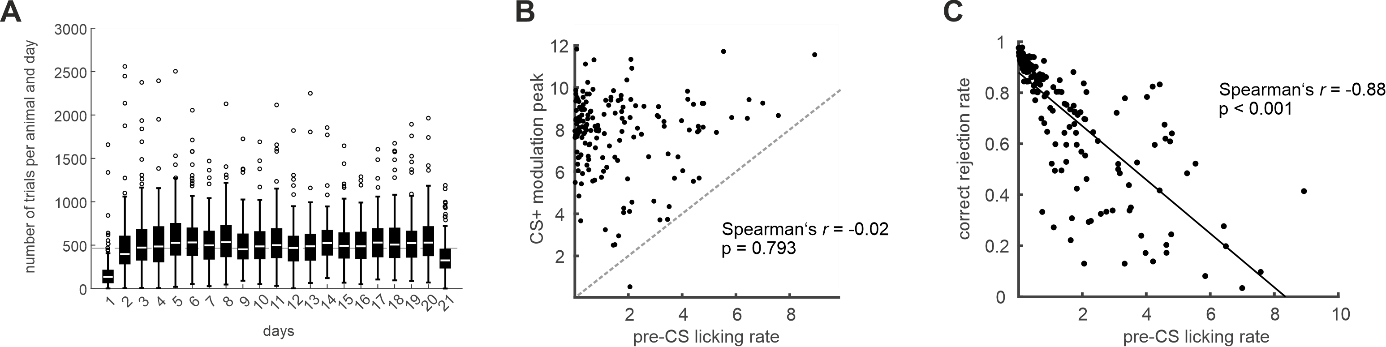

**Fig. S2: Number of trials at the stimulus-outcome (S-O) learning module and associations between reward-seeking features.**

**A,** Boxplots (median, 25th and 75th percentile) show the number of trials performed per animal per day at the S-O learning module during the first three weeks in the NoSeMaze. Whiskers extend to 1.5 x the interquartile range; individual points represent outliers. The data were obtained from 17 groups (each n = 9-10 male mice on C57BL/6J background).

**B,** No significant correlation was found between the pre-CS licking rate and the CS+ modulation peak. The CS+ modulation peak values were calculated from the 150 trials before each reversal (stable lick phase) across the 17 groups, while the pre-CS licking rate was computed from all trials. The dashed-line marks the diagonal.

**C,** The pre-CS lick rate and the correct rejection rate were strongly negatively correlated. All trials were used to calculate these rates.

*CS, conditioned stimulus*

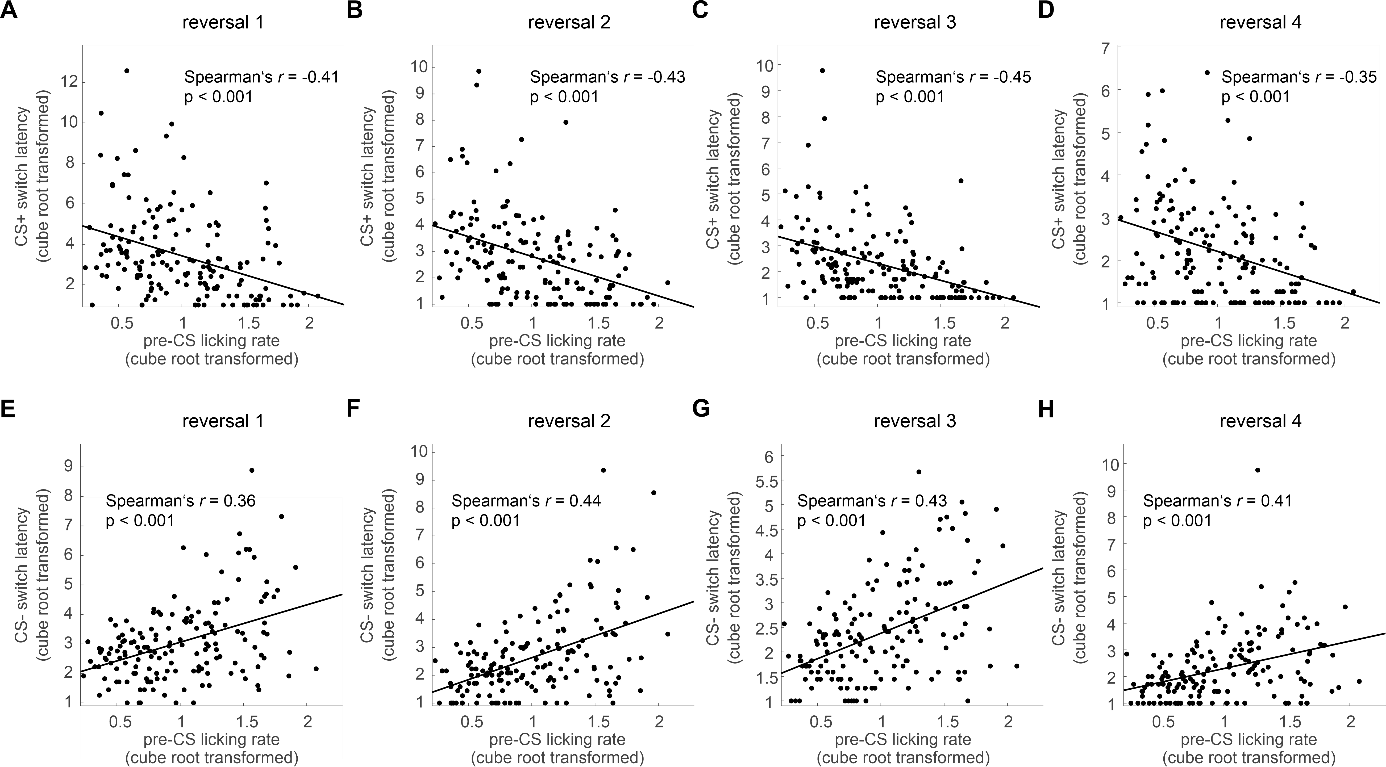

**Fig. S3: Association between pre-CS licking rate and switch latencies during reversal learning at the stimulus–outcome (S–O) learning module.**

**A–D**, The pre-conditioned stimulus (pre-CS) licking rate was negatively correlated with the switch latencies at the CS+ across reversals 1 to 4. Both variables were cube root transformed.

**E–H**, The pre-CS licking rate was positively correlated with the switch latencies at the CS− across reversals 1 to 4. Both variables were cube root transformed.

*CS, conditioned stimulus.*

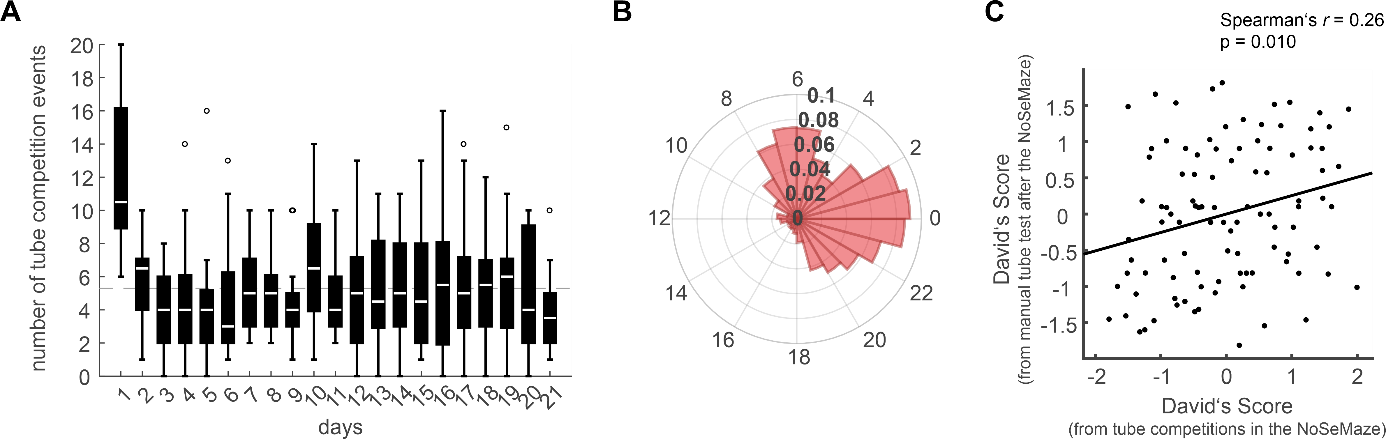

**Fig. S4: Number and timing of tube competitions and association between the David’s scores from manual tube tests and tube competitions in the NoSeMaze.**

**A,** Boxplots (median, 25th and 75th percentile) illustrate the number of tube competitions per day and group during the first three weeks in the NoSeMaze. The whiskers extend to the most extreme data points not considered outliers. Outliers are defined by 1.5 interquartile ranges. The mean value over all days is represented by a dashed line. The data is derived from 18 different groups (each n=9-10 male mice).

**B,** The polar histogram shows the distribution of the relative number of tube competitions across the 24-hour cycle. Most competitions occurred during the dark phase (between 20:00 and 08:00).

**C,** The David’s scores from tube competitions in the NoSeMaze over three weeks positively correlated with the David’s scores from manual tube tests after the animals had been removed from the NoSeMaze. The data were derived from 10 groups (each with 9-10 animals).

*NoSeMaze, non-invasive sensor rich maze*

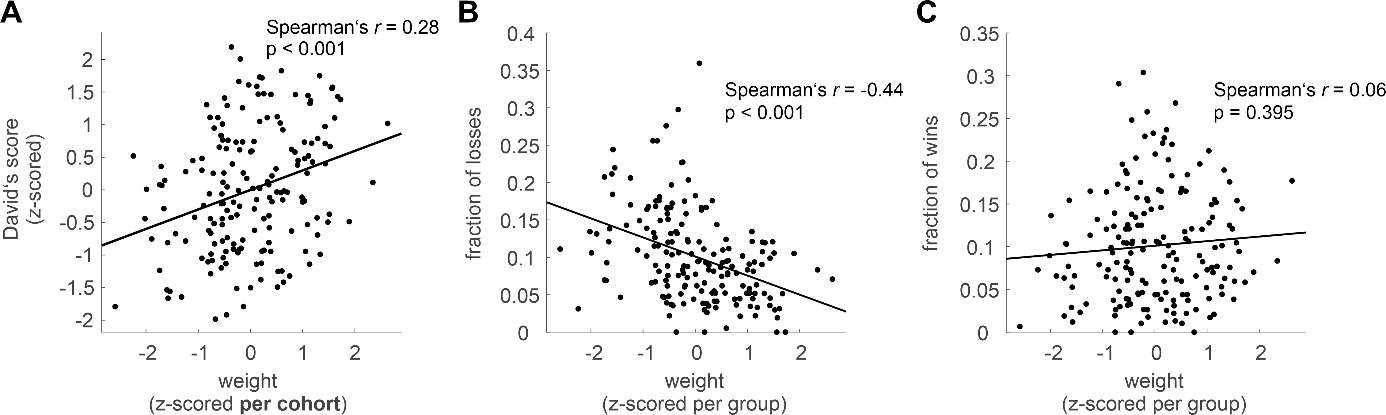

**Fig. S5: Relationship between body weight and outcomes of tube competitions in the NoSeMaze.**

**A**, Relative body weight before entering the NoSeMaze was positively associated with the David’s score.

**B**, Body weight (z-scored per group) was significantly negatively correlated to the fraction of losses in tube competitions.

**C**, No association was found between body weight and the fraction of wins in tube competitions.

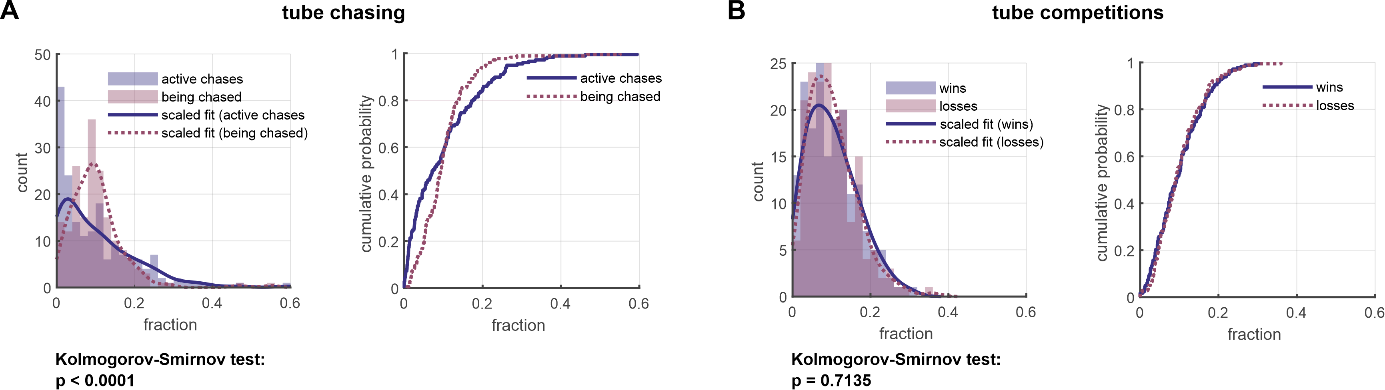

**Fig. S6: Distributions of win-loss outcomes from chasing and tube competitions in the NoSeMaze.**

**A,** Histograms show the fraction of active chases (violet) and times being chased (dark-red) in chasing events per animal, together with their scaled gamma fits (left panel). The two distributions were significantly different (Kolmogorow-Smirnov test, p < 0.0001). The cumulative probability plot of the same data is shown in the right panel, highlighting the differences between the distributions.

**B,** Histograms show the fraction of wins (violet) and losses (dark-red) in tube competitions per animal, together with their scaled fits (left panel). The two distributions did not differ significantly (Kolmogorow-Smirnov test, p = 0.0001). The cumulative probability plot of the same data is shown in the right panel, further illustrating the similarity between the distributions.

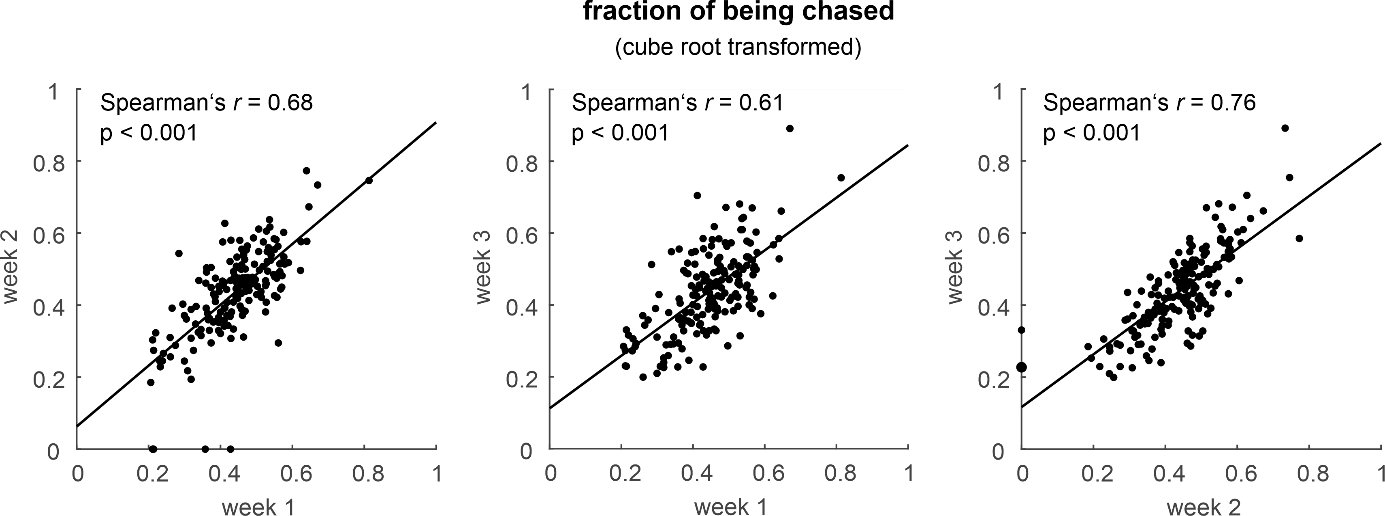

**Fig. S7: Association of the fraction of times being chased across different weeks in the NoSeMaze.**

**Scatterplots show that the fraction of times being chased was significantly correlated across all three weekly comparisons within one NoSeMaze round, indicating temporal stability of this metric. Data were obtained from 18 groups and cube-root transformed.**

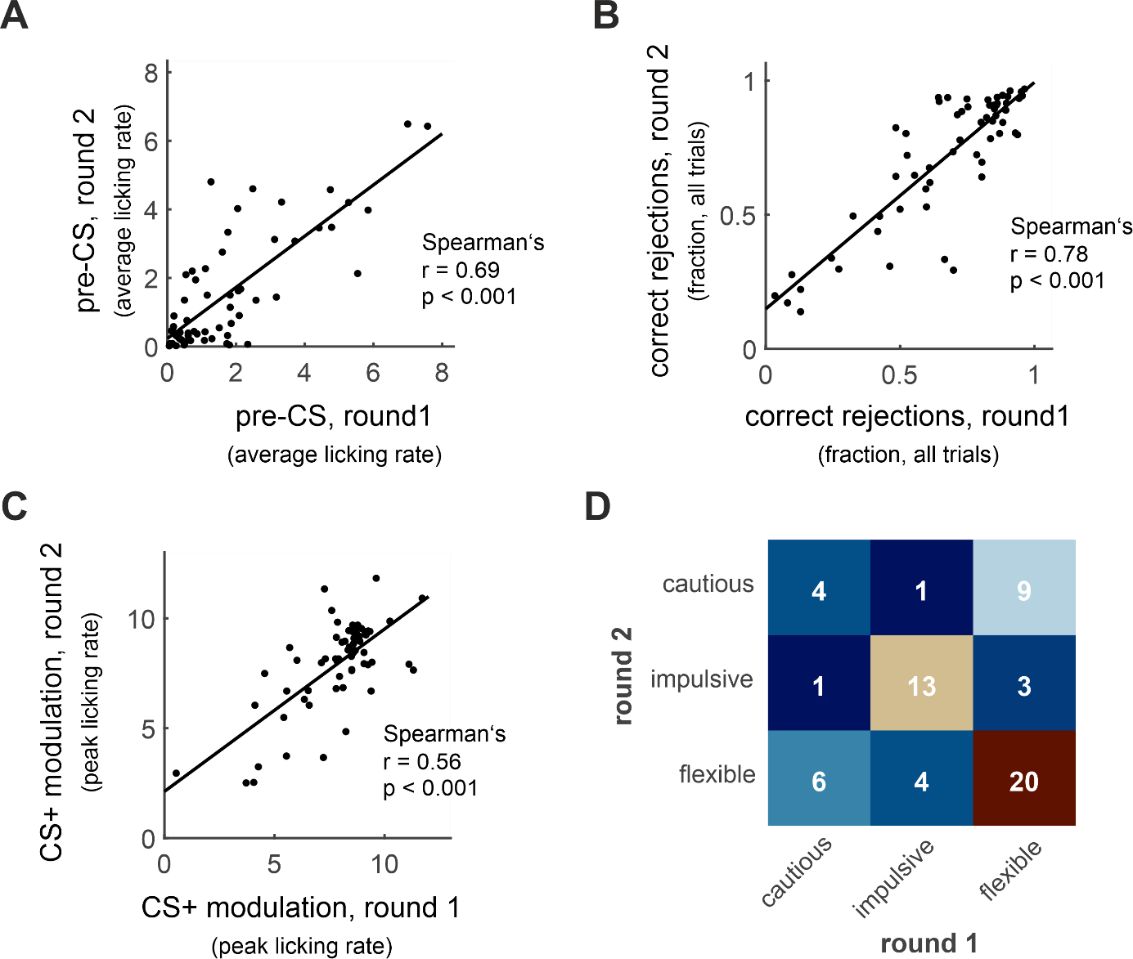

**Fig. S8: Stability of reward-seeking features across NoSeMaze rounds with different group members.**
**A–C,** Scatterplots show strongly significant positive correlations of individual behavioral metrics between round 1 and 2 in the NoSeMaze for (**A**) the average pre-CS licking rate, (**B**) the correct rejection rate, and (**C**) the CS+ modulation peak. Each dot represents one animal. Group compositions differed between the two rounds.
**D,** The consistency matrix shows individual behavioral clustering into cautious, impulsive, and flexible learners across both NoSeMaze rounds. Most animals retained their classification between rounds, indicating stability of behavioral learning profiles despite changes in group composition.

|  | **Observed median difference (round 1 - round 2)** | **p-value  (two-tailed permutation test on the median, n=10,000)** |
| --- | --- | --- |
| **CS+ switch latency** |  |  |
| **reversal 1** | 21 | 0.0008 |
| **reversal 2** | 10 | 0.0066 |
| **reversal 3** | 2.5 | 0.1140 |
| **reversal 4** | 2.5 | 0.0539 |
| **median (all reversals)** | 3.75 | 0.0122 |
| **CS- switch latency** |  |  |
| **reversal 1** | 28.5 | <0.0001 |
| **reversal 2** | 5 | 0.0626 |
| **reversal 3** | 6 | 0.0088 |
| **reversal 4** | 3 | 0.2706 |
| **median (all reversals)** | 7 | 0.0039 |

**Table S3: Differences in CS+ and CS- switch latencies between NoSeMaze rounds.**

Observed median differences (round 1 – round 2) are shown for each reversal and the overall median across all reversals. P-values were calculated using two-tailed paired permutation tests on the median (n = 10,000 permutations).

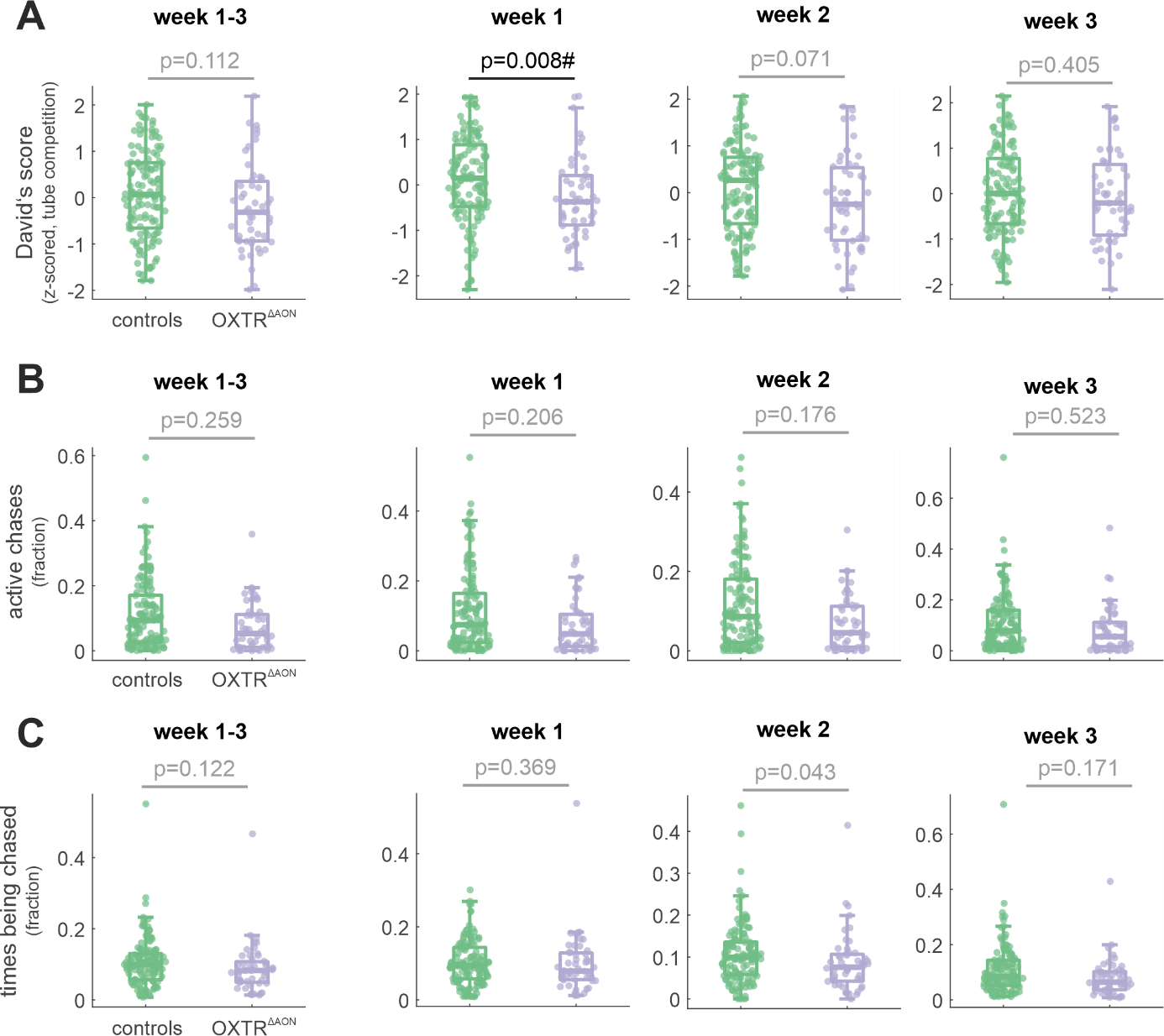

**Fig. S9: Effects of OXTR^ΔAON^ on social rank and chasing.**

**A–C**, Comparisons between control and OXTR^ΔAON^ mice for (**A**) David’s score (z-scored from tube competitions), (**B**) fraction of active chases, and (**C**) fraction of times being chased across all NoSeMaze rounds. The four panels show the comparison including all weeks (far left), and separately for week 1 (middle left), week 2 (middle right), and week 3 (far right). No significant group differences were detected, except for a transient reduction in the David’s score for week 1 (**A**, second panel). Group differences were assessed using permutation tests (n = 10,000) based on group medians. To account for repeated measures, group labels were shuffled at the level of the animal. Week-wise comparisons were marked with # if surviving FDR correction. The findings indicate that selective OXTR deletion in the AON pars centralis exerted only minor and short-lived effects on social rank and chasing.

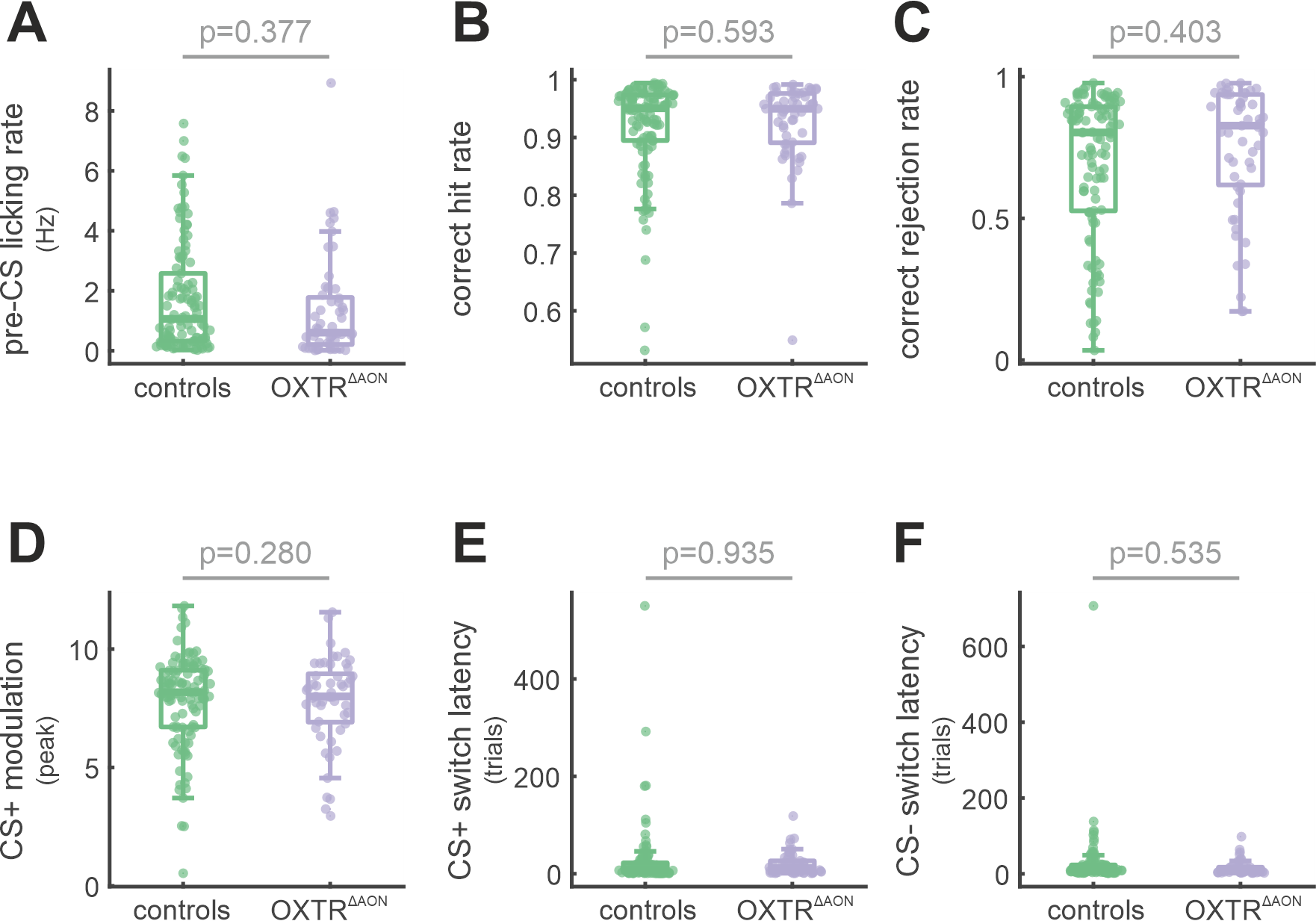

**Fig. S10: Effects of OXTR^ΔAON^ on reinforcement-learning measures in the NoSeMaze.**

**A–F**, Comparisons between control and OXTR^ΔAON^ mice for key reinforcement-learning features: (**A**) pre-CS licking rate, (**B**) correct hit rate, (**C**) correct rejection rate, (**D**) CS+ modulation peak, (**E**) CS+ switch latency, and (**F**) CS– switch latency. No significant group differences were detected in any measure. Group differences were assessed using permutation tests (n = 10,000) based on group medians, with group labels shuffled at the animal level to account for repeated measures. These results indicate that selective OXTR deletion in the AON pars centralis did not affect reinforcement-learning performance in the NoSeMaze.

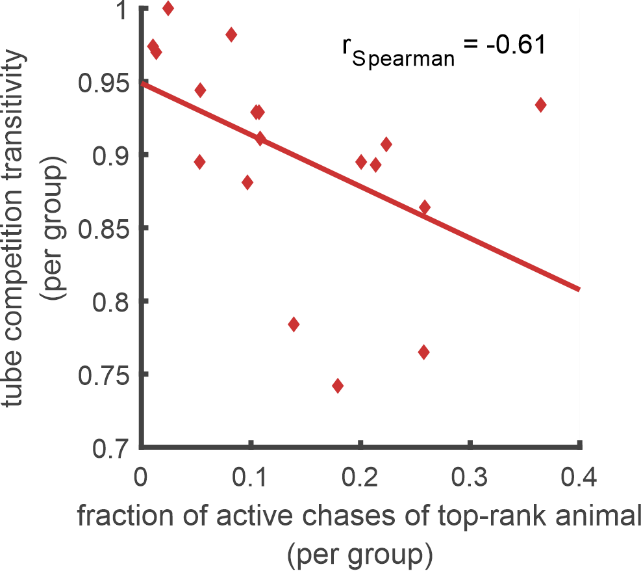

**Fig. S11: Association between active chases of the animal with the highest social rank and the transitivity of the social hierarchy in the respective group.**

Across groups, the fraction of active chases from the mouse with the highest social rank (ranking based on David’s scores from tube competitions) was negatively associated with the group-level transitivity. Groups with lower transitivity showed a higher fraction of active chases, suggesting that aggressive status signaling via active chases becomes more relevant when social hierarchies are less well-defined.

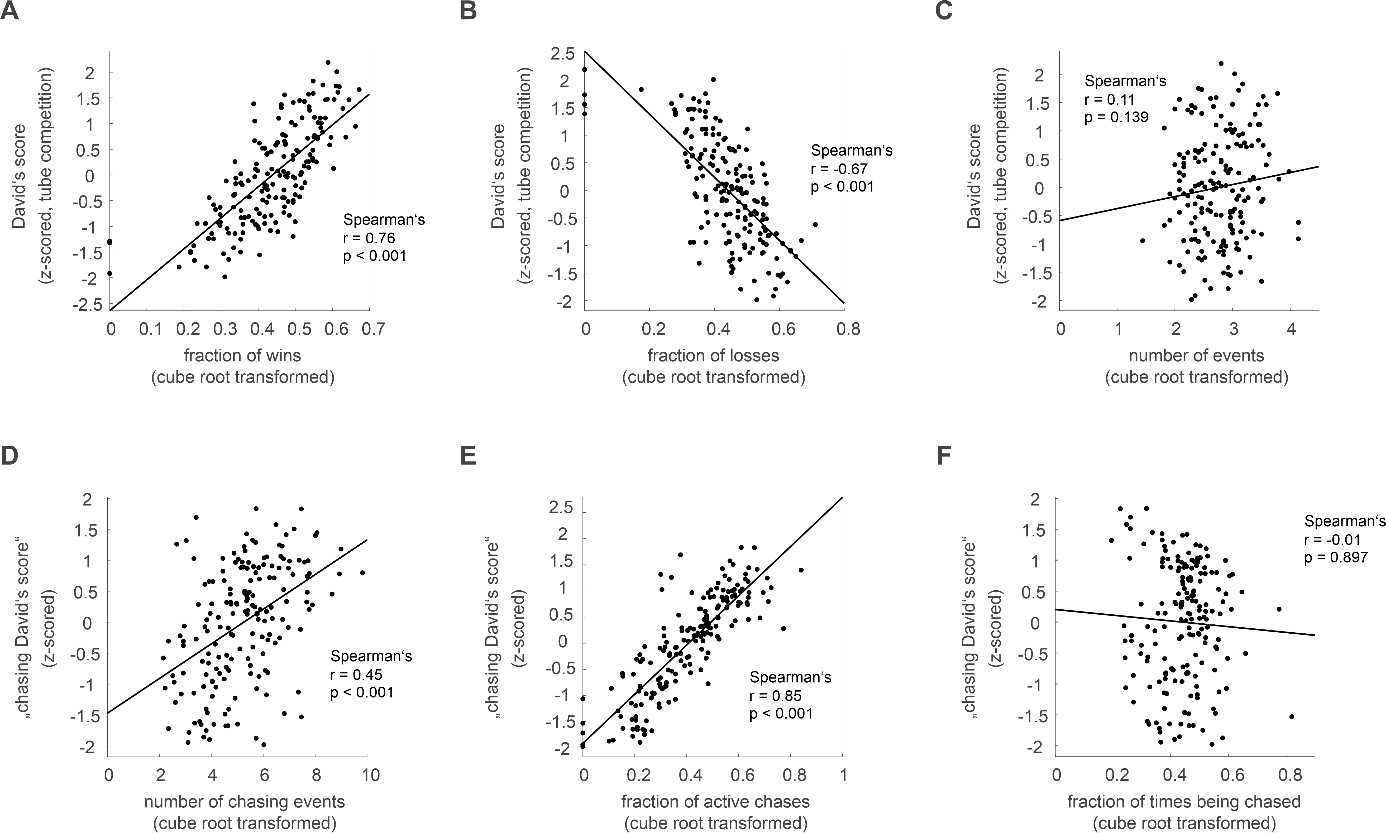

**Fig. S12: Validation of social rank metrics in tube competitions and chasing.**

**A–C,** Correlations between the z-scored David’s score (derived from automated tube competition data) and (**A**) the fraction of wins, (**B**) the fraction of losses, and (**C**) the total number of tube competition events per animal (all cube root transformed). David’s score was strongly associated with both the fraction of wins (r = 0.76) and the fraction of losses (r = –0.67), confirming symmetry in dominance relationships. No significant association was found between David’s score and competition frequency (r = 0.11, p = 0.139), suggesting that tube encounters occurred incidentally and were not strategically avoided.

**D–F,** Correlations between the z-scored “chasing David’s score” and (**D**) the number of chasing events, (**E**) the fraction of active chases, and (**F**) the fraction of times being chased (all cube root transformed). The “chasing David’s score” was positively associated with both the number of chases (r = 0.45) and the fraction of active chases (r = 0.85), but showed no relationship with the fraction of times being chased (r = –0.01). These findings indicate that chasing behavior was asymmetrically distributed across individuals and reflected volitional initiation rather than reciprocal dominance-subordination interactions.

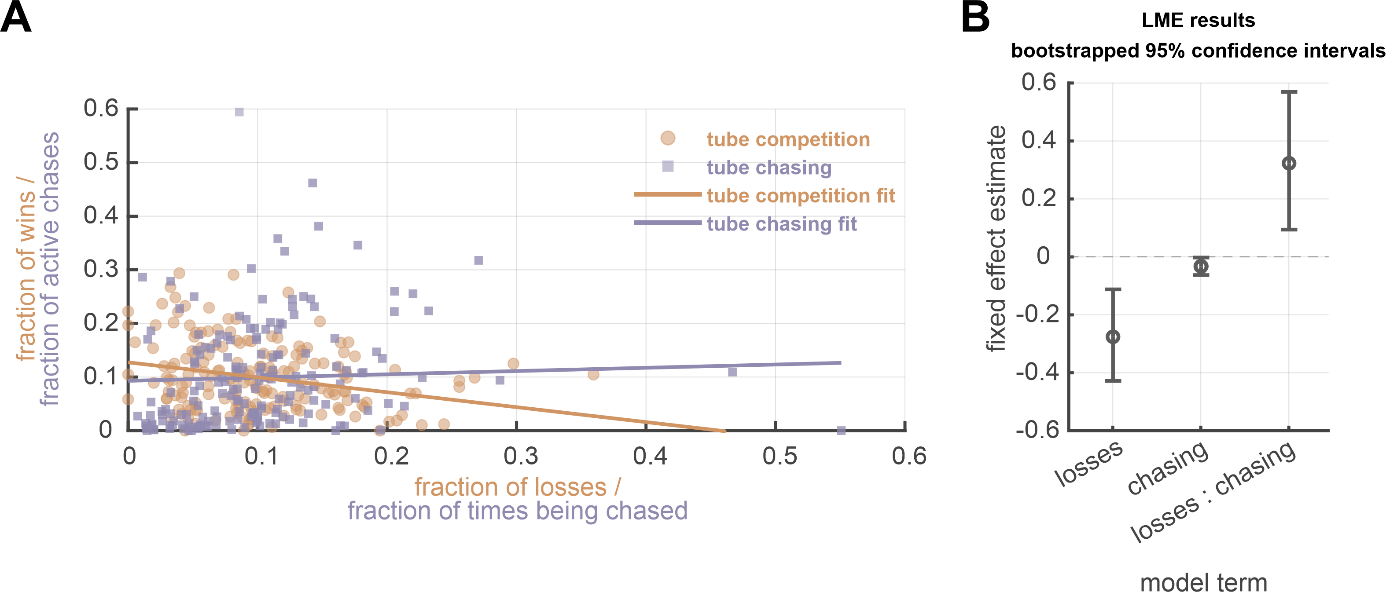

**Fig. S13: Divergent win–loss relationships in tube competitions and chasing.**

**A,** Scatterplot comparing the fraction of wins (orange) and the fraction of active chases (violet) against the corresponding fraction of losses and times being chased, respectively. Linear fits show that the strong negative relationship between wins and losses in tube competitions differs markedly from the flatter and mildly positive relationship observed for active chases and being chased.

**B,** Fixed-effect estimates from a linear mixed-effects model (LME) predicting the fraction of ‘wins’ across both behavioral modalities. The model included the fraction of ‘losses’, event type (tube competition vs. chasing), and their interaction as fixed effects, with mouse identity as a random intercept. Bootstrapped confidence intervals (10,000 cluster-resampled iterations) are shown for each term. A significant interaction confirmed that the relationship between win and loss fractions differs between tube competitions and chasing behavior, supporting a lack of symmetry in chasing.

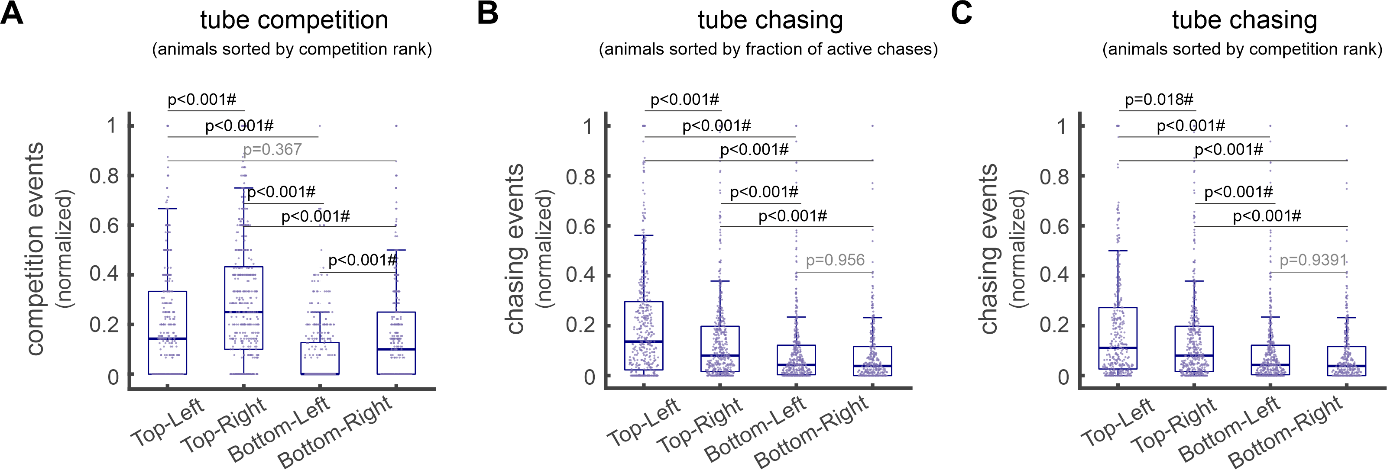

**Fig. S14: Distribution of normalized dyadic competition and chasing events across matrix quadrants.**

**A**, Boxplots show the distribution of normalized dyadic competition events counts for each quadrant of the competition matrix (cf. **Fig. 6D** in the main), where animals are sorted by their competition-based social ranks. The matrices were divided into four quadrants: top-left (dominant vs. dominant), top-right (dominant vs. subordinate), bottom-left (subordinate vs. dominant), and bottom-right (subordinate vs. subordinate). Each point represents a single dyadic value (i.e., a specific pair of individuals) from one group. Normalization was performed within each group by its maximum value to ensure comparability across groups.

**B**, Same as a, but for chasing event counts. Dyadic chasing events were extracted from matrices sorted by the fraction of active chases and analyzed by quadrant.

**C**, Same as b, but chasing events were extracted from matrices sorted by competition-based social ranks. Notably, the highest frequency of chasing events occurred in the top-left quadrant, indicating frequent chasings between individuals with high social ranks.

Beta LMEs were applied using quadrant as a fixed effect and group as a random intercept. Pairwise post hoc comparisons between quadrants were conducted on the response scale with Tukey-adjusted p-values for multiple comparisons (marked with #). For details, see Methods.

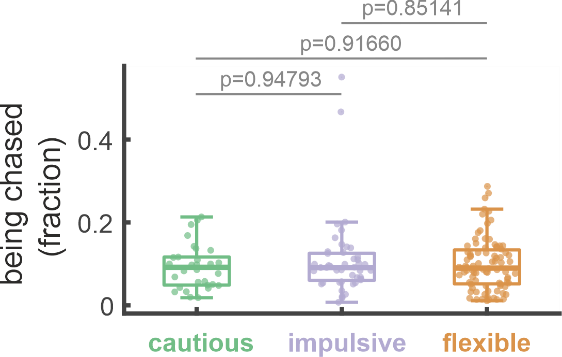

**Fig. S15: Comparison of the fraction of times being chased across the behavioral clusters defined from the reward-seeking task.**

No significant differences were found between the cautious, impulsive, and flexible clusters (p-values from permutation tests with n = 100.000 permutations using the median). Boxplots show the median (line), interquartile range (IQR, box), and whiskers extending to 1.5 × the IQR.
